# Mitotic errors drive rapid clearance of polyploidy during intestinal regeneration despite robust centrosome clustering

**DOI:** 10.64898/2026.03.25.714196

**Authors:** Iva Dundović, Kruno Vukušić, Thomas van Ravesteyn, Beatriz Carvalho, Marianna Trakala, Geert J.P.L. Kops, Iva M. Tolić

## Abstract

Polyploid cells are increasingly recognized not only as hallmarks of cancer but also as features of regenerating tissues. During intestinal regeneration, polyploid cells are transient, yet the mechanisms underlying their clearance remain unknown. Using mouse intestinal organoids as a regeneration model, we show that, unlike in many cancer cell-line models, this clearance occurs without immediate cell-cycle arrest and is not driven by failure to establish spindle bipolarity. Instead, polyploid intestinal cells efficiently cluster supernumerary centrosomes to form bipolar spindles in an HSET-dependent manner, facilitated by delayed centrosome separation at mitotic onset. Despite this, polyploid divisions frequently produce chromosome segregation errors, including catastrophic chronocrisis. Lineage tracing reveals that progeny of such divisions is rapidly lost over subsequent generations. Increasing polyploidy during early regeneration disrupts organoid maturation, indicating that timely polyploidy clearance is required for successful regeneration. Polyploid cells are also detected in regenerating human colonic organoids, suggesting that transient polyploidy is a conserved feature of intestinal regeneration.

## Introduction

Polyploidy, or whole genome doubling, is a state in which a cell possesses more than two complete chromosome sets. As one of hallmarks of tumorigenesis, whole genome doubling has been studied extensively in the context of tumour initiation, progression and evolution^1–5^. In tumor and pretumor cells, whole genome doubling usually results from unprogrammed cytokinesis failure, mitotic-exit failure, or mitotic-entry failure^6,7^. While cytokinesis failure usually leads to the formation of binuclear polyploids, other mechanisms tend to generate mononuclear polyploids^8^. Regardless of the route, whole genome doubling results in the accumulation of both extra DNA and centrosomes, which restrict cell proliferation through activation of p53-dependent pathways^1,9–11^. First, newly formed polyploids typically exhibit reduced proliferative capacity due to replication stress associated with increased DNA content, and activation of pathways that recognize extra centrosomes. Second, even when mitosis is permitted, polyploids usually assemble highly error prone multipolar spindles, which results frequently in rapid cell death. To mitigate this, polyploids cluster extra centrosomes to assemble less error-prone pseudo-bipolar spindles^12,13^, and by elevated replication stress associated with increased DNA content^8^. Furthermore, asynchronous cell cycle progression of different nuclei of a polyploid cell (“chronocrisis”), generates DNA damage at mitotic entry^14,15^. As a result of such processes, frequent chromosome missegregation and catastrophic karyotypes that emerge from polyploid divisions are important drivers of punctuated karyotype evolution in colorectal cancer^16^.

Despite clear oncogenic association, polyploidy is the normal physiological state for specific cell types. Physiologically scheduled polyploidy generally does not contribute to chromosome missegregation, as newly formed polyploids are typically non-proliferative and fulfill specialized functions, as observed in megakaryocytes, cardiomyocytes and trophoblast giant cells^6,17^. Regenerating liver is a notable exception exhibiting a high proportion of proliferative polyploid hepatocytes that are the result of programmed cytokinesis failure^18,19^. In adult hepatocytes undergoing regeneration, polyploid mitoses are predominantly bipolar with low chromosome segregation errors when tissue architecture is preserved^20^. Nevertheless, polyploid hepatocytes in chronic regeneration can undergo reductive mitosis that produce lower-ploidy progeny, a phenomenon described by Duncan et al.^21^ and confirmed through in vivo lineage tracing^22–24^. Dividing endopolyploid cells with mitotic errors have also been reported in Drosophila rectal papillary cells^25^.

More recently, polyploid cells have also been described in organoids derived from healthy mouse small intestine (mSI), although in a drastically smaller fraction than in regenerating liver^26^. These polyploid cells originate from programmed cytokinesis failure as a consequence of a regenerative state that arises during early organoid growth^27,26^. During regeneration, the developmental state of the intestinal cells is switched back to fetal-like signature, changing the expression of whole sets of genes^28–30^. One of the affected genes is the large tumor suppressor kinase 1 (LATS1), which gets downregulated during regeneration and in disease, preventing cytokinesis and leading to cell polyploidization^26,31^. Polyploid cells are extruded into the lumen or become part of short-lived villus as the mSI organoid exits regenerative state^26^, but mechanisms driving this clearance are unknown.

In this study, we tested several models of polyploidy evolution to identify key factors limiting polyploid propagation during intestinal regenerative state. We used regrowth of mSI cystic organoids to model early intestinal regeneration and combined it with live cell imaging, acute polyploidy induction, motor protein and microtubule polymerization inhibition, long-term live-cell progeny tracking and organoid formation assays. We found that in regenerating mSI cystic organoids, polyploid cells with extra centrosomes are enriched, yet they are progressively lost over time. This occurs even though polyploid cells efficiently and rapidly cluster their supernumerary centrosomes. Asymmetric clustering, a mechanism that can rapidly restore normal centrosome numbers, also takes place but does not support long-term propagation of polyploid cells. Thus, stable polyploid populations are not maintained, in contrast to cultured cell lines, and we found no evidence of depolyploidization via multipolar spindle divisions, as occurs in regenerating liver. Instead, polyploid progeny is selectively eliminated due to highly error-prone bipolar divisions, including catastrophic chronocrisis events in the first polyploid division. These mitotic errors arise despite intact apicobasal polarity. Enrichment of polyploidy in early organoids impairs their progression into organized, mature organoids, highlighting the importance of timely polyploidy clearance. Notably, we also observed polyploid cells in regenerating human cystic organoids of the colon. Together, our study provides the first detailed view of mitosis in regenerating intestinal polyploid cells, revealing the novel selective mechanisms that restrict polyploid persistence in a highly proliferative tissue context.

## Results

### Polyploidy is present in early intestinal organoids but progressively eliminated despite efficient clustering of extra centrosomes

We set out to explore polyploid proliferating cells in the early mouse small intestinal (mSI) cystic organoids^26^ and their fate as the organoid matures. We considered several models for polyploidy evolution that act through mitosis, basing them on the currently available knowledge in other systems (Figure 1A). In the first model, polyploid cells are allowed to continuously propagate and form a stable polyploid population (Figure 1A, model 1). This is possible if there is no penalty on the extra centrosomes^9^. To test this model in mSI regeneration, we chose two timepoints after seeding cells dissociated from mSI organoids, 48 and 96h, to enrich for either early, small or later, large cystic organoids, respectively (see Methods). To determine the fraction of dividing cells with more than two centrosomes, which we used as an indicator of acute polyploidization^13,33^, we imaged anaphase cells in fixed mSI cystic organoids with expression of EGFP-Centrin-2 and stained with DAPI (Figure 1B). We additionally stained cells for α-tubulin and F-actin to confirm that centrin foci colocalize with tubulin foci to make functional spindle poles and that they are within cell bounds (Supplementary Fig. 1A). The fraction of mitotic cells with more than two centrosomes (which we term *proliferating cells with extra centrosomes*, PCEC) accounted for 16% of divisions in small organoids (Figure 1B, Supplementary Fig. 1A). The fraction of PCEC quickly decreased during organoid growth as 6% of PCEC remained in middle-sized organoids and only 2% in large organoids. The increase in metaphase cell size and spindle morphometric measures^34–37^ of PCEC, compared to cells with two centrosomes (Supplementary Fig. 1B), further suggests that PCEC are indeed polyploids. Interestingly, at 48h after dissociation, all of the mitotic single cells from the mSI culture were PCEC, with the majority containing more than 4 centrosomes (80%, Supplementary Fig. 1C), likely reflecting limited ability of single PCEC to form larger organoids due to repetitive cytokinesis failure. These results suggest that PCEC are present in early mSI cystic organoids that are undergoing regeneration but are selected against as the organoid grows and matures, which argues against the model of continued proliferation of polyploids while maintaining extra centrosomes (Figure 1A, model 1, left).

**Figure 1.**
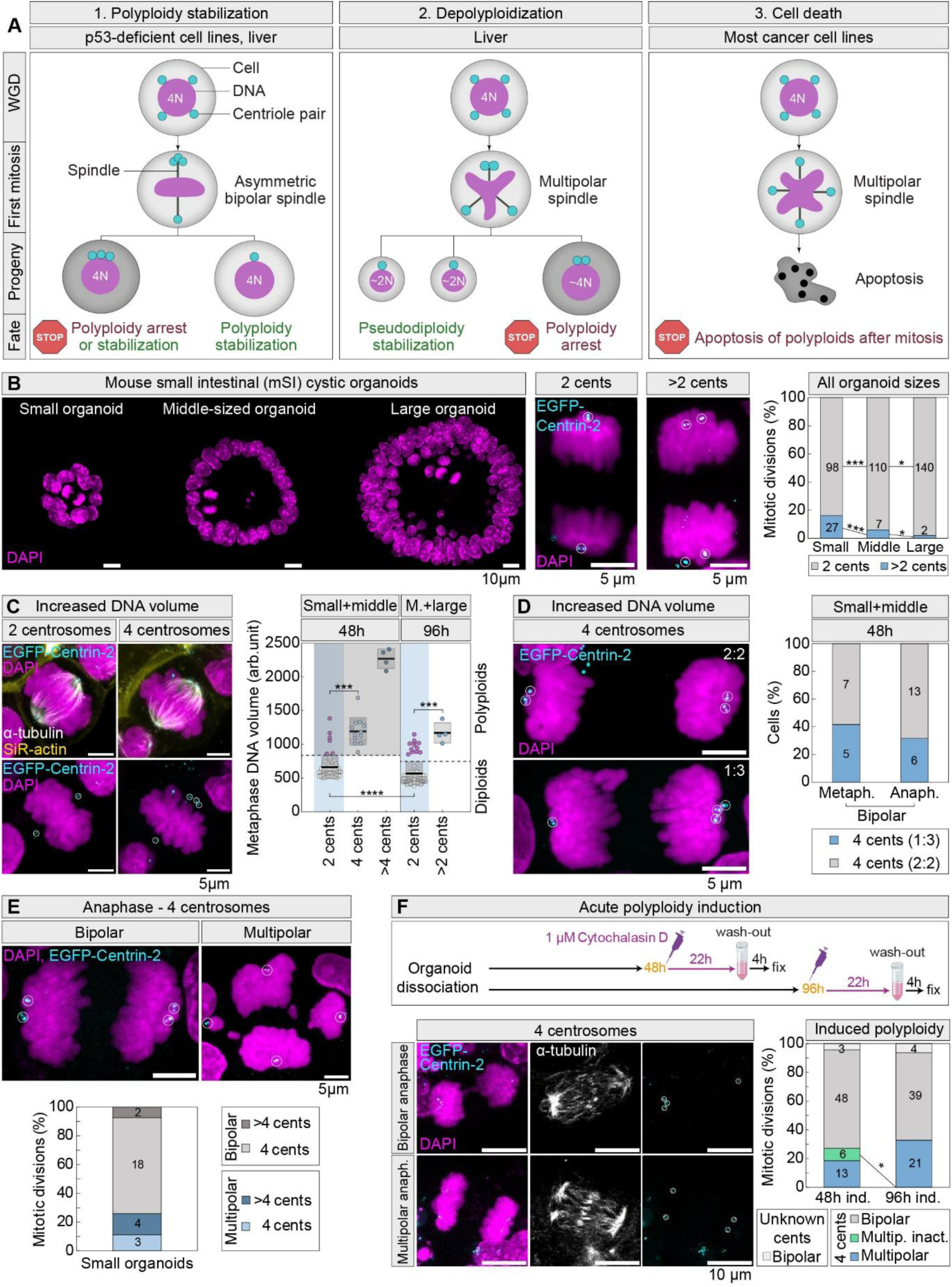
Loss of extra centrosomes or their efficient clustering is not sufficient to prevent the elimination of polyploid cells in mouse small intestinal (mSI) early organoids. (**A**) Current mitotic models of polyploidy evolution in various systems, based on existing literature (see main text). STOP signs represent cell cycle arrest or cell death. Note that polyploid cells with extra centrosomes can be arrested prior to the first mitosis (not shown) or after the first mitosis by p53-dependent pathways. (**B**) Fixed images of mSI organoids (left) and anaphase cells (middle) stably expressing EGFP-Centrin-2 (cyan) and stained with DAPI (magenta). The bar plot shows the frequencies of anaphase cells with two or more than two centrosomes in organoids of different sizes. N = 384 cells from > 300 organoids and ≥ 3 experiments. (**C**) Fixed images of mSI metaphase cells stably expressing EGFP-Centrin-2 (cyan), immunostained for α-tubulin (grey) and stained for F-actin with SiR-actin (yellow) and for DNA with DAPI (magenta). The univariate scatter plot shows metaphase DNA volume, obtained with segmentation analysis, in cells with different centrosome numbers. Colored points represent individual cells; black lines show the mean, with light and dark grey areas marking 95% confidence intervals for the mean and standard deviation, respectively. Cells are separated by pseudotime of growth, which is based on the time of fixation post seeding (48h and 96h) and the size of the organoid from which they were taken. Dashed lines indicate the minimum threshold for increased DNA volume, associated with polyploidy, defined as the mean + SD of cells with two centrosomes (see Methods). Blue shaded areas indicate cells with two centrosomes and the grey shaded area indicates cells with increased DNA content at 48h. N = 157 cells from > 100 organoids and ≥ 3 experiments. (**D**) Images show fixed bipolar mSI anaphase cells with four centrosomes clustered in a symmetric (2:2) or asymmetric (1:3) configuration. The frequency of each configuration in metaphase and anaphase is shown on the graph on the right, at pseudotime described in (**C**). N = (12, 19) cells from > 10 organoids and ≥ 3 experiments; p > 0.05. (**E**) Fixed mSI anaphase cells with more than two centrosomes exhibiting bipolar or multipolar spindles (top panel) and frequency of each type in small organoids (bottom panel). N = 27 cells from > 20 organoids and ≥ 3 experiments. (**F**) Top panel: schematic representation of the protocol for acute polyploidy induction with cytochalasin D. Numbers in yellow represent timepoints after seeding of dissociated organoids; numbers in purple time of incubation with the inhibitor and numbers in black incubation time with fresh medium before fixation (recovery time). Images on the bottom show fixed mSI anaphases after polyploidy induction. Graph shows quantification of spindle polarity types in anaphase after polyploidy induction at either 48 or 96h post seeding dissociated organoids. N = (70, 64) cells from > 60 organoids and ≥ 3 experiments. All images are maximum intensity projections. White circles mark centriole pairs for which centrosome identity was confirmed using α-tubulin staining (tubulin channel not always shown). Statistics: Z-test for two population proportions when comparing frequencies in (B), (D) and (F); in (C) Kruskal-Wallis test with post hoc Dunn’s test. Symbols: *, P < 0.05; ***, P ≤ 0.001; ****, P ≤ 0.0001. For simplicity, in (B) and (C) not all p-value symbols are shown. Cents, centrosomes; arb., arbitrary; M., middle; metaph., metaphase; anaph., anaphase; fix, fixation; inact., inactivated; multip., multipolar; ind., induced.

Loss of extra centrosomes could be a rapid strategy to survive harmful polyploidy, as previously observed^13,32^. We detected dividing polyploids with only two centrosomes by performing chromosome segmentation analysis of metaphase cells in mSI cystic organoids and measuring their metaphase plate volume, which correlates strongly with their ploidy (Figure 1C, Supplementary Fig. 1D), similarly to previous observations^34–37^. This approach allowed us to estimate the proportion of cells with increased DNA content but normal centrosome counts (Figure 1C, dashed threshold, see Methods). We found that ∼10% (n=6 out of 57) of cells with two centrosomes at 48h exhibited DNA content indistinguishable from that of PCEC (Figure 1C, purple points within blue area), indicating polyploidy. Those cells accounted for ∼27% of all cells with increased DNA content at 48h after dissociation (n=6 out of 22, purple points within grey area), suggesting that PCEC are losing centrosomes as a result of a single cell division event. A similar fraction of cells with possible centrosome loss was observed in larger organoids at 96h (15%, Figure 1C, purple points within blue area), although the total fraction of PCEC decreased (∼22% at 48h to ∼5% at 96h, Figure 1C) consistent with measurements in anaphase (Figure 1B). Taken together, polyploids accounted for 30% (22 out of 73) of divisions in organoids at 48h, and 19% (16 out of 84) at 96h, indicating a substantial reduction of polyploidy over just 2 days of organoid maturation.

In order for PCEC to lose centrosomes in one generation, bipolar spindles should be able to cluster their centrosomes in an asymmetric 1:3 configuration^32^. Indeed, we found that ∼30% of PCEC clustered their anaphase centrosomes in an asymmetric configuration in our system (Figure 1D), consistent with previous reports in 2D models of freshly generated polyploids^32^ and with the fraction of cells with suspected centrosome loss at 48h (Figure 1C). A similar fraction of cells with asymmetric clustering was observed in metaphase (Figure 1D). These results suggest that the rapid loss of extra centrosomes resulting from asymmetric clustering (Figure 1A, model 1) contributes to the adaptation of early mSI cystic organoids to polyploidy, although it is not sufficient to sustain their propagation to larger cystic organoids.

These findings directed us to test additional models, in which polyploidy is eliminated through frequent multipolar divisions that either produce viable near-diploid progeny (Figure 1A, model 2)^21–23^ or cease to divide due to poor viability^13^ (Figure 1A, model 3). Multipolar divisions in early mSI cystic organoids accounted for less than 30% of total PCEC divisions (Figure 1E, 25.9% in small organoids, n=7 out of 27) and less than 15% (14.3%, n=3 out of 21) when considering only PCEC with four centrosomes. This contrasts with the poor ability of non-cancer epithelial cell lines to cluster extra centrosomes^13,32,38^, unless E-cadherin is depleted^39^. Since E-cadherin was present in mSI cystic organoids (Supplementary Fig. 1E), this discrepancy likely reflects the differences associated with tissue-of-origin.

Given the high prevalence of bipolar divisions in PCEC (Figure 1E), we hypothesized that the prevalence of bipolar divisions could be driven by a negative selection that prevents proliferation of the highly error-prone multipolar cells that were inefficient in clustering^40^. To test this, we acutely induced polyploidization by blocking cytokinesis with cytochalasin D^13^ for 22h, followed by a drug washout and fixation of cystic organoids 4h later (Figure 1F). Polyploidy was induced both 48h after dissociation, where PCEC are more abundant, and after 96h, where they are rare (Figure 1F and 1C). We found that a majority of freshly induced polyploids with four centrosomes formed a bipolar spindle (Figure 1F, 68.6% and 60.9% when induced at 48h or 96h respectively), while only about 30% of cells divided with some form of a multipolar spindle (Figure 1F, 27.1% and 32.8% when induced at 48h or 96h respectively). The fraction of multipolar divisions was similar to untreated small organoids (Figure 1E) and the trend remained the same when excluding the population of multipolar spindles with partial or total pole inactivation (8.6%, Supplementary Fig. 1F), defined as spindles with a distant pole containing weak tubulin signal and without adjacent chromosomes. Additionally, symmetric centrosome clustering was prevalent in induced PCEC bipolar spindles (Supplementary Fig. 1G), similar to spindles in uninduced PCEC (Figure 2B, right). We conclude that efficient centrosome clustering in early mSI organoids is an inherent property of mSI cells, which is reminiscent of highly efficient clustering of polyploid hepatocytes^20,21,33^.

**Figure 2.**
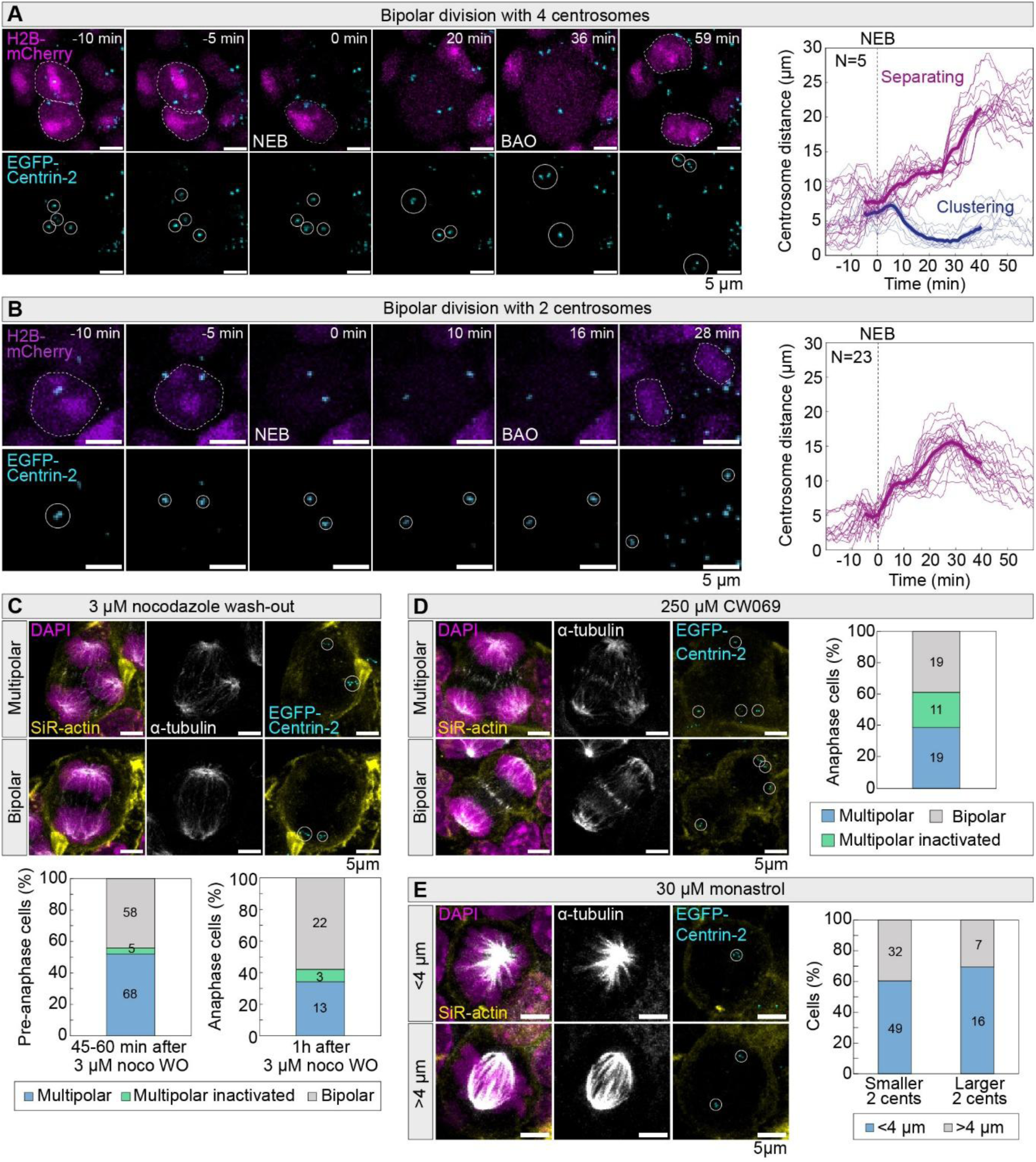
Rapid and efficient clustering of extra centrosomes in early mSI organoids is driven by delayed centrosome separation and unbalanced motor activity. **(A)** Representative time-lapse images of a bipolar division in a mSI cell with four centrosomes stably expressing EGFP-Centrin-2 (cyan) and with leaky expression of H2B-mCherry (magenta). Plot shows distances between all tracked centrosomes in cells with four centrosomes over time. Centrosome distances that are < 6 µm apart before anaphase onset are defined as clustering (blue thin lines) and otherwise as separating (purple thin lines). N = 5 cells from > 3 organoids and ≥ 3 experiments. **(B)** Time-lapse images of a bipolar division in a mSI cell with two centrosomes. Plot shows centrosome distances in time (purple thin lines) for all cells with two centrosomes that had diploid-sized nuclei (see Methods). N = 23 cells from > 10 organoids and ≥ 3 experiments. In (A-B) time zero marks mitotic onset (dashed line); thick lines are mean values; shaded areas are SEM values. **(C)** Fixed images of mSI anaphase cells stably expressing EGFP-Centrin-2 (cyan), immunostained for α-tubulin (grey) and stained for F-actin with SiR-actin (yellow) and for DNA with DAPI (magenta). Cells are fixed 45-60 min post wash-out of 3 µM nocodazole. The bar plots show quantification of mitotic spindle polarity after nocodazole wash-out, in cells prior to (left) and during (right) anaphase. N = (131, 38) cells from (> 100, > 30) organoids and ≥ 3 experiments. **(D)** Fixed images of mSI anaphase cells after 30 min of treatment with 250 µM HSET inhibitor CW069. In (C-D), prior to treatment, polyploidy was induced using cytochalasin D at 48h post seeding dissociated organoids and only cells with four centrosomes were selected for analysis. The bar plots show quantification of mitotic spindle polarity in treated cells. N = 49 cells from > 40 organoids and ≥ 3 experiments. **(E)** Fixed images of mSI mitotic cells with two centrosomes after 3h of treatment with 30 µM Eg5 inhibitor monastrol. The bar plot shows the frequency of centrosome distances <4 µm or >4 µm, with larger cells shown separately due to possible polyploidy (see Methods). N = (81, 23) cells from (> 50, > 15) organoids and ≥ 3 experiments. Statistics: Z-test for two population proportions when comparing frequencies in (E). All images are maximum intensity projections. White circles mark centriole pairs for which centrosome identity was confirmed using α-tubulin staining or by observing centriole pair trajectories during live-cell imaging (see Methods). The cell nuclei and chromatin groups in telophase are outlined with white dashed lines for clarity. The cell nuclei and chromatin groups in telophase are outlined with white dashed lines for clarity. NEB, nuclear envelope breakdown; BAO, before anaphase onset; noco WO, nocodazole wash-out; cents, centrosomes.

Cumulatively, our results show that polyploid cells with extra centrosomes are a feature of early mSI organoids and are selected against during organoid growth. The low tendency of PCEC to undergo multipolar divisions due to highly efficient clustering suggests that both depolyploidization and elimination of progeny of multipolar divisions (Figure 1A, model 2 and 3), are not major drivers of polyploidy clearance. Furthermore, the asymmetric clustering that leads to rapid centrosome loss (Figure 1A, model 1) is not sufficient to stabilize a population of polyploids in later organoid stages suggesting that another mechanism may determine the fate of polyploid cells.

### Rapid and efficient clustering of extra centrosomes in early organoids is enforced by delayed centrosome separation

Polyploids within mSI cystic organoids showed an innate property to cluster their extra centrosomes very efficiently, leading to distinct pathways of polyploidy evolution in these organoids in comparison to other cell systems. Efficient clustering of extra centrosomes depends on the activity of motor proteins^12,41–43^ whose efficiency of clustering is modulated by centrosome distances at the onset of mitosis^39^. To explore positioning of extra centrosomes in mSI organoids in time, we live-imaged small cystic organoids expressing EGFP–Centrin-2 48h after seeding. The leaky H2B-mCherry expression was used to visualize nuclei, identify nuclear envelope breakdown (NEB) and telophase chromosome groups (Figure 2A). Tracking of centrosome distances of natural PCEC in time showed that at NEB centrosome pairs that end up separating in anaphase were on average 7.6 ± 1.6 µm apart (Figure 2A), compared to 4.96 ± 1.52 µm in cells with two centrosomes (Figure 2B). Distances between centrosomes in mSI were similar in mature organoids (Supplementary Fig. 2A). These distances were substantially lower than in 2D-grown diploid and polyploid epithelial cell lines^39^, which typically separate centrosomes to larger distances, reflecting movement to opposing sides of the nucleus prior to NEB (prophase spindle assembly pathway)^39,44^. The distance between separating centrosomes in PCEC increased toward metaphase (12.8 ± 0.6 µm) and peaked in anaphase (24.5 ± 2.9 µm), whereas the distances in cells with two centrosomes were consistently lower (Figure 2A-B). A short centrosome distance before NEB was tied to the formation of the prometaphase rosette in some cell lines (prometaphase spindle assembly pathway)^45–48^. In agreement with short distances prior to NEB, fixed prometaphase cells in untreated mSI cystic organoids all assembled prometaphase rosettes (Supplementary Fig. 2D-E). We conclude that polyploid cells within the early epithelial mSI organoids, unlike epithelial cell lines^39^, dominantly use the prometaphase pathway of spindle assembly independently of their ploidy.

To find out whether this late centrosome separation promotes extra centrosome clustering in PCEC, we tested the correlation between initial centrosome pair distance and its ability to cluster in mitosis within individual live-imaged cells. Average centrosome distance at NEB in PCEC was similar for centrosome pairs that clustered (6.2 ± 2.9 µm) and those that separated before anaphase (7.6 ± 1.6 µm) (p=0.087, Figure 2A, Supplementary Fig. 2F, left). We conclude that centrosome distance at NEB is not a strong predictor of clustering efficiency within mSI cells, although the shorter centrosome spacing observed in our system compared to others, could facilitate more efficient clustering^39,49^.

To further test whether clustering success is constrained by the initial spatial arrangement of centrosomes in mSI cystic organoids, we performed a nocodazole washout experiment. Before washout, mean initial centrosome distance in induced polyploids was 6.8 ± 4.1 µm (Supplementary Fig. 2G), similar to live imaging measurements of PCEC (Supplementary Fig. 2F, left). After nocodazole washout, only a third (∼34%) of cells remained multipolar in anaphase (Figure 2C), confirming that clustering in regenerating organoids is robust, at least in part due to naturally restricted centrosome spacing at mitotic entry.

Since the minus end–directed motors are essential for pole clustering in other systems^12,41–43,50^ and as their activity is promoted by the short initial centrosome distancing^39^, we hypothesized that they also contribute to this process in mSI PCEC. Indeed, inhibition of the major minus-end directed motor HSET with CW069^51^ caused a dose-dependent increase in multipolar metaphases, reaching nearly 50% at 250 μM and ∼80% at 400 μM (Supplementary Fig. 2H). In anaphase, ∼39% of cells remained multipolar in anaphase at 200 μM CW069, whereas anaphases at 400 μM were rare (Figure 2D). Clustering efficiency could be also promoted by the decreased activity of outward spindle forces^42,43^. Indeed, mSI PCEC were highly sensitive to partial inhibition of the outward force-generating motor Eg5, as treatment with 30 μM monastrol reduced centrosome spacing to <4 μm in ∼60% of cells (Figure 2E). This sensitivity contrasts with non-tetraploid mammalian cells, which require substantially higher monastrol concentrations to induce spindle collapse^42,52^. Together, these findings indicate that mSI polyploid cells operate close to the threshold of outward Eg5-dependent force generation, thereby biasing spindle mechanics toward HSET-driven inward clustering.

Finally, we quantified the speed of clustering in mSI PCEC. Using the time from NEB to anaphase onset as a proxy for clustering dynamics^12,38,51^, we observed that mitotic duration in mSI PCEC increased with centrosome numbers, from 17.8 ± 4.9 minutes in cells with two centrosomes to 29 ± 4.5 minutes in cells with four (p= 0.002, Supplementary Fig. 2F, right). Despite this increase, mSI polyploid cells clustered supernumerary centrosomes within 20-30 minutes, which is ∼2 times faster than in freshly generated polyploid HCT-116 or RPE-1 cells, where clustering takes ∼50 minutes^38,53^. In agreement, the clustering process started almost immediately after NEB in PCEC (Figure 2A), contrary to cell lines^39^. Together, these data support a model in which delayed centrosome separation, restricted initial spacing, and a motor-force balance close to the outward-force threshold promote both robust and unusually rapid clustering of extra centrosomes in mSI cystic organoids.

### Polyploid cells in early organoids are prone to anaphase errors, partially due to frequent chronocrisis

Proliferating polyploids in different cellular contexts either drastically decrease mitotic fidelity and survival^1,13^ or are well tolerated and enable proficient mitosis^20,26^. As we have shown that mSI polyploids dominantly divide in a bipolar way, but are still cleared from the population, contrary to most polyploid systems, we hypothesized that PCEC bipolar divisions are error-prone. Analysis of all dividing cells in mSI cystic organoids (Figure 3A) revealed that anaphase and telophase cells had a low rate of chromosome missegregation (Supplementary Fig. 3A), independent of the organoid size (12.4% in small, 7% in middle-sized and 7.9% in large organoids). However, a significant portion of PCEC produced an error in anaphase (46.9%, n=15 out of 32 within small and middle-sized organoids cumulatively) (Figure 3A, left bar plot), indicating that PCEC do not divide faithfully. PCEC erroneous divisions contributed to roughly half of the total segregation errors in small and middle-sized organoids (Figure 3A, right bar plot, 55.6% and 50% respectively, purple shades). Bipolar divisions that make up the majority of PCEC divisions (Figure 1D, 74.1%, n=20, N=27, Supplementary Fig. 3B) showed a high tendency to missegregate chromosomes (Figure 3B, 35% error rate, n=7, N=20), proving to be the main contributor to mitotic error rate in mSI organoids. We conclude that polyploid cells that arise during intestinal regeneration are highly error prone during mitosis, despite highly efficient clustering of extra centrosomes. Mitotic errors were rare in cells with two centrosomes (5.7% in small, 4.5% in middle-sized, and 6.1% in large organoids), but accounted for a substantial proportion of total errors (44.4%, 50%, and 85.7%, respectively; Figure 3A). Segmentation analysis showed that most of these cells did not exhibit DNA content consistent with PCEC (Supplementary Fig. 3C–D), indicating that mitotic errors in cells with two centrosomes are not linked to prior polyploidization.

**Figure 3.**
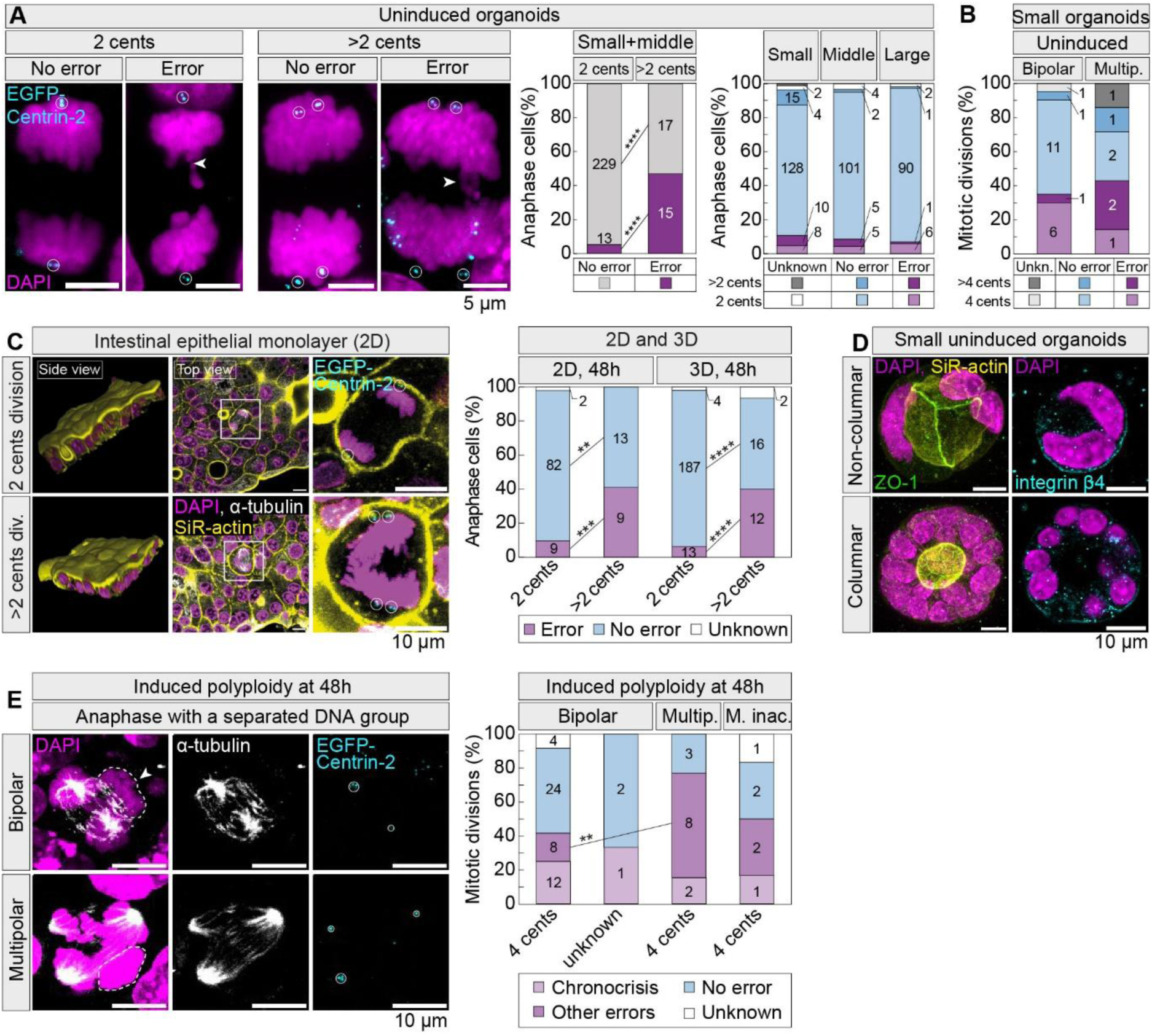
Polyploid cells in early mSI organoids are prone to anaphase errors, in part due to frequent chronocrisis. **(A)** Fixed images of mSI anaphase cells with two or more than two centrosomes, stably expressing EGFP-Centrin-2 (cyan), immunostained for α-tubulin (not shown) and stained for F-actin with SiR-actin (not shown) and for DNA with DAPI (magenta). The bar plot shows the frequency of anaphase segregation errors, depending on centrosome number (left) or organoid size (right). Cells from large organoids are excluded from the left plot due to a low number of cells with more than two centrosomes. N = (242, 32) cells from (> 200, >30) organoids and ≥ 3 experiments. Organoid sizes reflect growth stages. Numbers of analysed cells are indicated on the data labels. Data from > 90 organoids and ≥ 3 experiments in each organoid group. **(B)** Frequencies of chromosome segregation errors in mSI anaphase cells with more than two centrosomes, depending on their spindle polarity. N = (20, 7) cells from (> 15, > 5) organoids and ≥ 3 experiments in each group. **(C)** Fixed images of mSI anaphase cells dividing within a monolayer, with two or more than two centrosomes. White rectangle marks the cell enlarged on the right. The first column shows the mSI monolayer as a three-dimensional projection. The bar plot shows the frequencies of chromosome segregation errors in mSI anaphase cells in a monolayer (“2D”) versus cells coming from organoids (“3D”) at the same timepoint after seeding dissociated organoids (48h). Numbers of analysed cells are indicated on the data labels. 3D data from > 30 organoids; ≥ 3 experiments in each group. **(D)** Images show fixed organoids immunostained for apicobasal polarity markers β4-integrin (cyan) and Zonula occludens-1 (ZO-1, green) and stained with SiR-actin (yellow) and DAPI (magenta). Images that show β4-integrin staining are single z-planes. **(E)** Fixed images of mSI anaphase cells after polyploidy induction with cytochalasin D that had separated chromatin groups lacking adjacent spindle poles, indicating chronocrisis. White arrows point to chromatin groups without adjacent poles, additionally encircled with white dashed lines. The bar plot shows the frequencies of chromosome segregation errors in anaphase cells after polyploidy induction at 48h post seeding dissociated organoids. Cells are separated in groups based on centrosome numbers and spindle polarity. Erroneous divisions coming from cells in chronocrisis are separated from other error types. Numbers of analysed cells are indicated on the data labels. Data from > 3 organoids and ≥ 3 experiments in each group. All images are maximum intensity projections, unless stated otherwise. “Uninduced” stands for native mSI organoids in which no polyploidy induction was performed, contrary to “induced polyploidy”. White circles mark centriole pairs for which centrosome identity was confirmed using α-tubulin staining (tubulin channel not always shown). Statistics: Z-test for two population proportions in (A)-(C), (E). Symbols: **, P ≤ 0.01; ***, P ≤ 0.001; ****, P ≤ 0.0001. For simplicity, in (C) not all p-value symbols are shown. Cents, centrosomes; unkn., unknown; div., division; multip. or m., multipolar; inac., inactivated.

To test if the chromosomal instability observed in PCEC depends on the organoid architecture, we transferred mSI cells from an organoid to a natural monolayer system, where cells grow in mostly 2D patches on a layer of extracellular matrix^54^. At 48h after seeding, PCEC could be detected in the population of dividing cells within the monolayer in a similar fraction as in mSI organoids at the same timepoint (Supplementary Fig. 3E-F). As in early mSI organoids (Figure 3A), the overall segregation errors were rare in the monolayer system, and PCEC accounted for most of the errors (Figure 3C). This shows that PCEC are inherently error-prone during mitosis, independent of their environment.

Chromosome segregation errors in early mSI organoids may result from impaired apicobasal polarity that preferentially affects polyploids, similar to what has been reported in immature primary mammary spheroids^20^. To test this possibility, we investigated epithelial polarity and morphology of early mSI organoids. Staining of small mSI organoids for polarity markers, including F-actin, Zonula occludens-1 (ZO-1), β-integrin and E-cadherin^20^, showed that apicobasal localization is established and clear lumina is formed (Figure 3D and Supplementary Fig. 4A-B)^55^. Additionally, we did not observe an increase of anaphase errors in immature organoids that did not reach the columnar organization of the organoid epithelium^56–58^, compared to those that did (Supplementary Fig. 4A-C), contrary to previous observations in mammary spheroids^20^. These results suggest that impaired mitotic fidelity observed in early mSI organoids is not the result of lost apicobasal polarity.

To identify processes driving the high mitotic error rate in mSI PCEC, while avoiding errors caused by accumulated DNA damage from past polyploid divisions, we analyzed mitotic errors in acutely induced polyploids (Figure 3E). In these cells, bipolar divisions with four centrosomes resulted in anaphase segregation error in 41.7% of cases, comparable to the ∼35% observed in PCEC of untreated small and middle-sized organoids (Figure 3E and 3B). Although lower than in multipolar divisions (∼77%), error rates of bipolar polyploids in mSI organoids are surprisingly high compared to freshly generated bipolar polyploid RPE1 cells (∼10% lagging chromosomes^13^) or bipolar polyploid hepatocytes in adult livers (∼5% lagging chromosomes^20^). This may be caused by a high proportion of chronocrisis^14,15^ in induced polyploid mSI cells (25%, Figure 3E and Supplementary Fig. 4D). Anaphase cells in chronocrisis had one chromosome group with markedly reduced DNA condensation that lacked adjacent spindle poles, probably due to asynchrony in mitotic entry of the two nuclei^14^ and those cells were predominantly associated with anaphase errors (Figure 3E, Supplementary Fig. 4D-F). Chronocrisis events were also observed in fixed, uninduced mSI PCEC, but at low frequencies (6.7%, Supplementary Fig. 4G), as expected given that in uninduced cells at 48h, a fraction of polyploids is mononucleated^26^. The high proportion of bipolar mitosis in mSI cystic organoids could be related to frequent chronocrisis events, as the amount of DNA interacting with the mitotic spindle is reduced during chronocrisis^59^. However, excluding cells in chronocrisis from the analysis of freshly induced polyploids in anaphase did not change the predominance of bipolar divisions (Supplementary Fig. 4H), indicating that chronocrisis is not a major driver of efficient centrosome clustering, though it does markedly increase anaphase error rates.

Altogether, our results show that the polyploid cells in early mSI organoids are highly error prone despite efficient clustering of centrosomes and preserved apicobasal polarity, in part due to frequent chronocrisis events in the first polyploid division.

### Mitotic errors drive rapid clearance of polyploids during organoid growth

To determine whether the frequent chromosome missegregation observed in mSI polyploids contributes to their clearance, we live-imaged mSI organoids expressing mNeonGreen–H2B for 2 days (Figure 4A) after induction of polyploidy with 1 μM cytochalasin D at 48 h post-seeding (as in Figure 1F), and compared them to DMSO-treated controls imaged under identical conditions. In cytochalasin D–treated samples, we tracked the progeny of binucleated cells, which we identify as polyploids (Figure 4A), whereas in DMSO-treated samples, random cells were tracked. We observed that the first divisions of polyploid cells were predominantly bipolar and frequently error-prone, consistent with both uninduced and induced fixed analyses of PCEC (Supplementary Fig. 5A; Figures 1 and 3). Both mother and daughter cells exhibited a low probability of forming multipolar spindles (Figure 4B), further confirming that efficient centrosome clustering is an intrinsic property of PCEC (Figure 1). Daughter cells arising from bipolar, error-free divisions typically underwent subsequent divisions (Figure 4C–D), indicating that polyploid progeny can proliferate when mitosis proceeds without errors. In contrast, approximately 24% of daughters of binucleated and bipolar mothers died during interphase, and this occurred almost exclusively when the mother cell had undergone an erroneous division (Figure 4C–D). Importantly, cell death following missegregation was also observed in DMSO-treated cells, despite the low frequency of erroneous divisions, whereas no cell death was observed in control diploid cells that divided without errors across two generations (Figure 4D), indicating that chromosome missegregation, rather than polyploidy per se, is the primary trigger of cell elimination. This effect persisted across generations, as granddaughter cells were more likely to die if mitotic errors had occurred in the first generation (Figure 4E), suggesting that the consequences of segregation errors extend beyond a single division. Surprisingly, daughters of erroneous multipolar divisions often retained proliferative capacity and underwent further division, although cell death was also increased across two generations (Figure 4C–D). The partial proliferative capacity of multipolar progeny may be explained by their frequent binucleation (Figure 4F), which has previously been associated with improved survival^13,60^. Cumulatively, these results indicate that propagation of PCEC in mSI organoids is primarily limited by frequent chromosome segregation errors, which drive rapid lineage-level elimination within a few generations.

**Figure 4.**
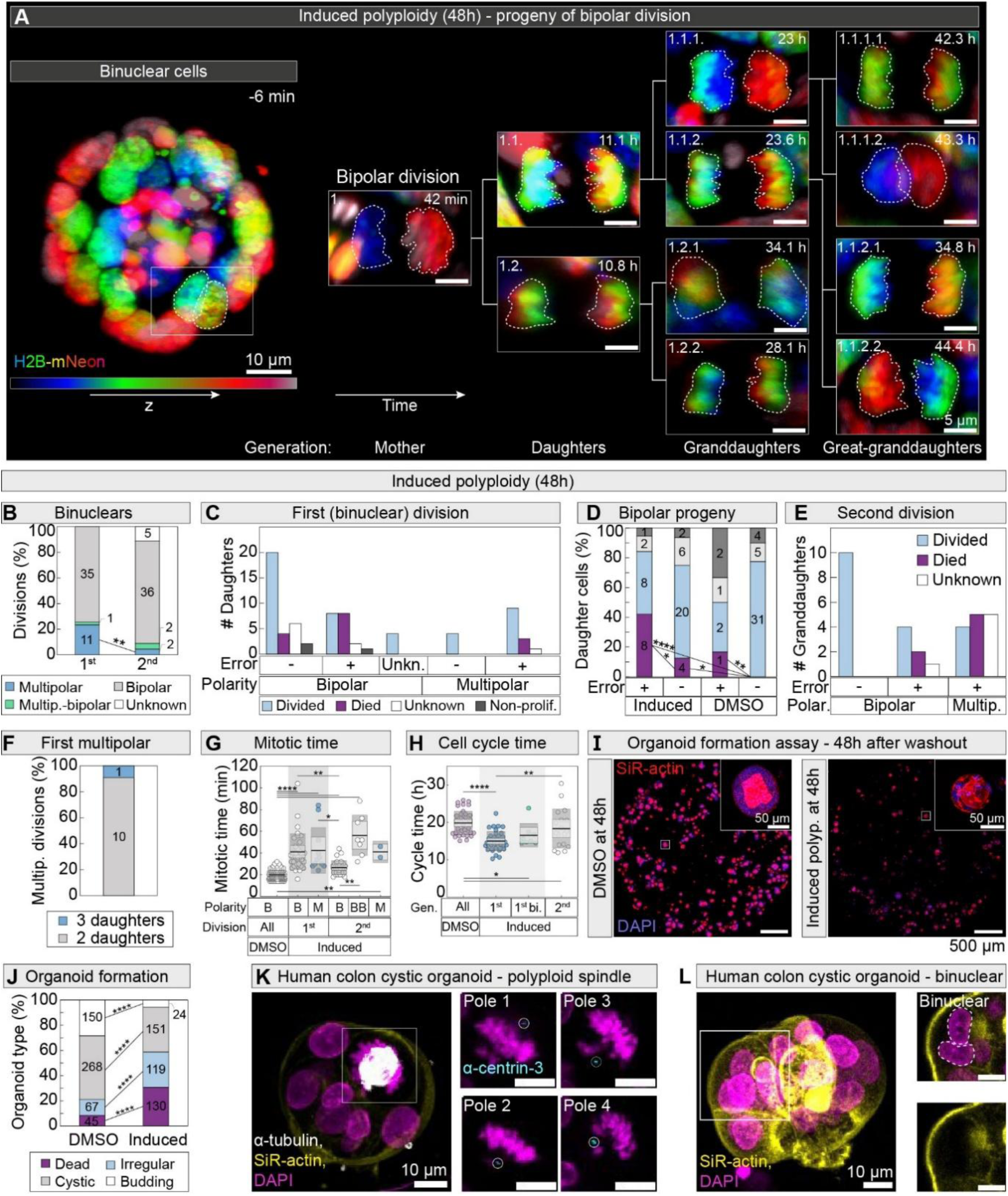
Proliferative polyploids are selectively eliminated from mSI organoids after mitotic errors and fail to generate mature organoids. **(A)** Representative time-lapse images of the progeny of a binuclear polyploid bipolar division in an mSI cystic organoid stably expressing H2B-mNeon (color coded by depth). Polyploidy was induced with cytochalasin D at 48h after seeding dissociated organoids. Binuclear polyploid cell is marked with a white rectangle. Time zero is mitotic onset. Numbers in left corners denote cell identity with respect to time and mother cell of origin. Nuclei of a binuclear cell and chromatin groups in anaphases are outlined with white dashed lines for clarity. Images of anaphase are smoothed with 0.5 sigma Gaussian blur. **(B)** The bar plot shows the frequencies of anaphase spindle polarity types in the first and second division after polyploidization, in live- imaged mSI cells expressing H2B-mNeon. N = (47, 45) cells from > 3 organoids and ≥ 3 experiments for each group. **(C)** The bar plot shows the number of the daughter cells undergoing denoted proliferation outcomes, based on their mother (first division) polarity status and presence of segregation errors during anaphase. Non-proliferative outcome is defined as daughter cells not dividing or dying for >30h after the mother division (see Methods). N = 72 cells from 3 organoids and 3 experiments. **(D)** The bar plot shows the frequencies of daughter cell proliferation outcomes depending on the segregation error status of the mother cell after polyploidy induction and in DMSO-treated controls. Legend as in (C). Number of cells is denoted on the data labels. Data from 3 organoids and 3 experiments. **(E)** The bar plot shows the frequency of multipolar divisions that resulted in either 2 or 3 daughter cells. Only first multipolar divisions after polyploidization were analysed. N = 11 cells from > 3 organoids and ≥ 3 experiments in total. **(F)** The bar plot shows the number of granddaughter cells undergoing denoted proliferation outcomes, separated based on spindle polarity type and chromosome segregation error outcome of the first (mother) division. All shown granddaughters come from bipolar daughters. N = 31 cells from 2 organoids and 2 experiments. **(G)** The univariate scatter plot of mitotic time for indicated spindle polarity types, shown separately for DMSO-treated control cells and cells with induced polyploidy. Shaded area (grey) separates cells in their first division after polyploidization from those in second division (right white area) and DMSO-treated control cells (left white area). N = (93, 34, 11, 30, 8, 2) cells from > 3 organoids and ≥ 3 experiments in total. **(H)** The univariate scatter plot of cycle duration for indicated generations. Cycle times of binucleated daughters are separated (“1^st^ bi.”). Data is shown separately for DMSO-treated control cells and cells with induced polyploidy. Shaded area (grey) separates cells in their first generation after polyploidization (daughter) from those in second generation (granddaughter, right white area) and DMSO-treated control cells (left white area). N = (37, 37, 8, 18) cells from > 3 organoids and ≥ 3 experiments in total. **(I)** Large-field images of fixed mSI organoids stably expressing EGFP-Centrin-2 (not shown), stained for F-actin with SiR-actin (red) and for DNA with DAPI (blue). Organoids were either treated with DMSO for control (left) or cytochalasin D to induce polyploidy (right), 48h after seeding dissociated organoids. Time of fixation is 48h after DMSO or inhibitor wash-out. Images in the right upper corners are enlargements of representative organoids marked with white rectangles. **(J)** Quantification of organoid types (see Methods) from (I). N = (530, 424) organoids from ≥ 3 experiments in total. **(K)** Fixed images of human non-cancer colonic organoids, immunostained for α-tubulin (grey) and centrin-3 (cyan), stained for F-actin with SiR-actin (yellow) and for DNA with DAPI (magenta). Single z-planes of a mitotic cell marked with a white rectangle are enlarged on the right to show centrosomes. White circles mark centriole pairs for which centrosome identity was confirmed using α-tubulin staining. **(L)** Fixed images of human non-cancer colonic organoids, stained as in (K), with single z-planes of a binuclear cell marked with a white rectangle enlarged on the right. Nuclei of a binuclear cell are outlined with white dashed lines. All images are maximum intensity projections, unless otherwise stated. In univariate plots, colored points represent individual cells; black lines show the mean, with light and dark grey areas marking 95% confidence intervals for the mean and standard deviation, respectively. Statistics: Z-test for two population proportions when comparing frequencies in (B), (D) and (J); in univariate plots: Kruskal-Wallis test with post hoc Dunn’s test. Symbols: *, P < 0.05; **, P ≤ 0.01; ****, P ≤ 0.0001. For simplicity, in (D) not all p-value symbols are shown. Multi or M, multipolar; non-prolif., non-proliferating; unkn., unknown; polar., polarity; gen., generation; B, bipolar; BB, binuclear bipolar; bi., binuclear; polyp., polyploidy; org., organoids.

### Mitosis is mildly extended in intestinal polyploid cells, whereas overall cell-cycle duration is unchanged

Polyploid progeny may also be eliminated during a longer time due to slower acting mechanisms. For example, increased mitotic duration of polyploids could activate mitotic stopwatch-based elimination^11,61^. After induced cytokinesis failure, the first polyploid mitosis was longer than the second mitosis or mitosis in uninduced PCEC, independently of polarity (Figure 4G and Supplementary Fig. 2F, right). Any type of polyploid division was still longer than diploid DMSO-treated controls (Figure 4G), and these differences are unlikely to result from mononucleation or asymmetric centrosome clustering, as previous work showed that neither phenomenon could cause observed changes in mitotic duration^53,62^. We conclude that increased mitotic duration could contribute to polyploid clearance in the long term.

In addition to mitosis-related effects, diploids may outcompete polyploids due to slow-acting fitness differences unrelated to mitosis, such as replication stress that leads to prolonged G1/S phase in polyploid cells^8^. To determine whether differences in overall cell-cycle length contribute to the loss of polyploid cells, we compared cell-cycle duration between induced polyploid cells and control cells. Surprisingly, no increase in cell-cycle duration was observed in polyploid cells and their progeny relative to controls (Figure 4H). As mitotic stopwatch activation is expected to increase G1 duration^63^, our results indicate that global cell-cycle prolongation, whether mitosis-dependent or independent, is unlikely to explain the progressive disappearance of polyploid cells during mSI regeneration.

### A transient burst of global polyploidization impairs organoid maturation

Our results thus far suggest that polyploid clearance could be important for normal organoid development. To assess whether organoids with predominantly polyploid cells can form mature, structured organoids over the long-term culturing, we devised a strategy to enrich polyploid cells during early regeneration. We induced polyploidy with cytochalasin D 48h after seeding dissociated organoids, and after further 48h of recovery following drug washout, we fixed the samples (similar to Figure 1F). In a parallel workflow, control samples were treated with DMSO instead of cytochalasin D. Fixed samples were then stained with F-actin to determine cell borders (Figure 4I and Supplementary Fig. 5B). Following cytochalasin D treatment, the formation of organoids with a clear epithelial structure was reduced, while dead structures increased. The living organoids were predominantly cystic and irregular, compared to more budding morphology observed in controls (Figure 4I-J). These results show that global polyploidization during early regeneration of intestinal organoids impairs their ability to form mature, organized intestinal organoids.

### Polyploidy is also detected in regenerating healthy human colon organoids

Finally, we asked whether polyploid cells we observed during regeneration of mSI cells are also present during the regenerative state of human gut cells^64^. Previously, polyploid cells were observed in samples from human colon cancer^16,65^, but evidence in healthy human colon organoids is lacking. Using dissociated organoids derived from normal tissue of a human donor, we established human colonic (hC) cystic organoids in the same manner as mSI cystic organoids (see Methods). Immunolabelling of Centrin-3 and α-tubulin confirmed presence of PCEC in small hC organoids fixed at 48h-post seeding (Figure 4K). Additionally, we found examples of binucleated interphase cells in fixed hC organoids of various sizes (Figure 4L). This suggests that polyploidy is a common feature of regenerating intestinal tissues with its clearance being an important feature of maintaining intestinal homeostasis across different species.

## Discussion

### Polyploidy elimination during regeneration in mSI

Early mSI organoid growth is an established regenerative-like model^27^ that enabled us to examine the fate of naturally occurring intestinal polyploids. Polyploid mSI cells frequently contained extra centrosomes but efficiently clustered them to form bipolar spindles before anaphase (Figure 1). About a third of polyploids can rapidly restore normal centrosome numbers through asymmetric clustering of extra centrosomes in mSI organoids (Figure 1), as reported in cell lines^32^. Although both efficient and asymmetric clustering could, in principle, stabilize a polyploid population^32^, we observe selective elimination of polyploid cells over longer time scales, largely driven by their high chromosome segregation error rates (Figures 1, 3 and 4). Mitotic errors in mSI organoids do not appear to result from loss of apicobasal polarity, contrary to mammary cells^20^, but instead arise from the presence of extra centrosomes or DNA early in growth (Figure 5, left path). Polyploid mSI cells are inherently error-prone and can undergo catastrophic chronocrisis events^15^ in their first division (Figure 5, right path). These findings point to a novel fast-acting mechanism that removes polyploid cells after erroneous divisions, preventing their further propagation during regeneration (Figure 5). Such rapid elimination could be particularly effective in the intestine, where proliferative windows are short and cell-cycle times are fast^66,67^. Consistent with this, we detect polyploids not only in mSI organoids but also in hC organoids, indicating that this phenomenon occurs in highly proliferative intestinal epithelia across both species and intestinal regions. Future work should define the molecular basis of this early error-detection pathway and determine whether it is sufficient to limit polyploid expansion or whether additional mechanisms act in parallel.

**Figure 5.**
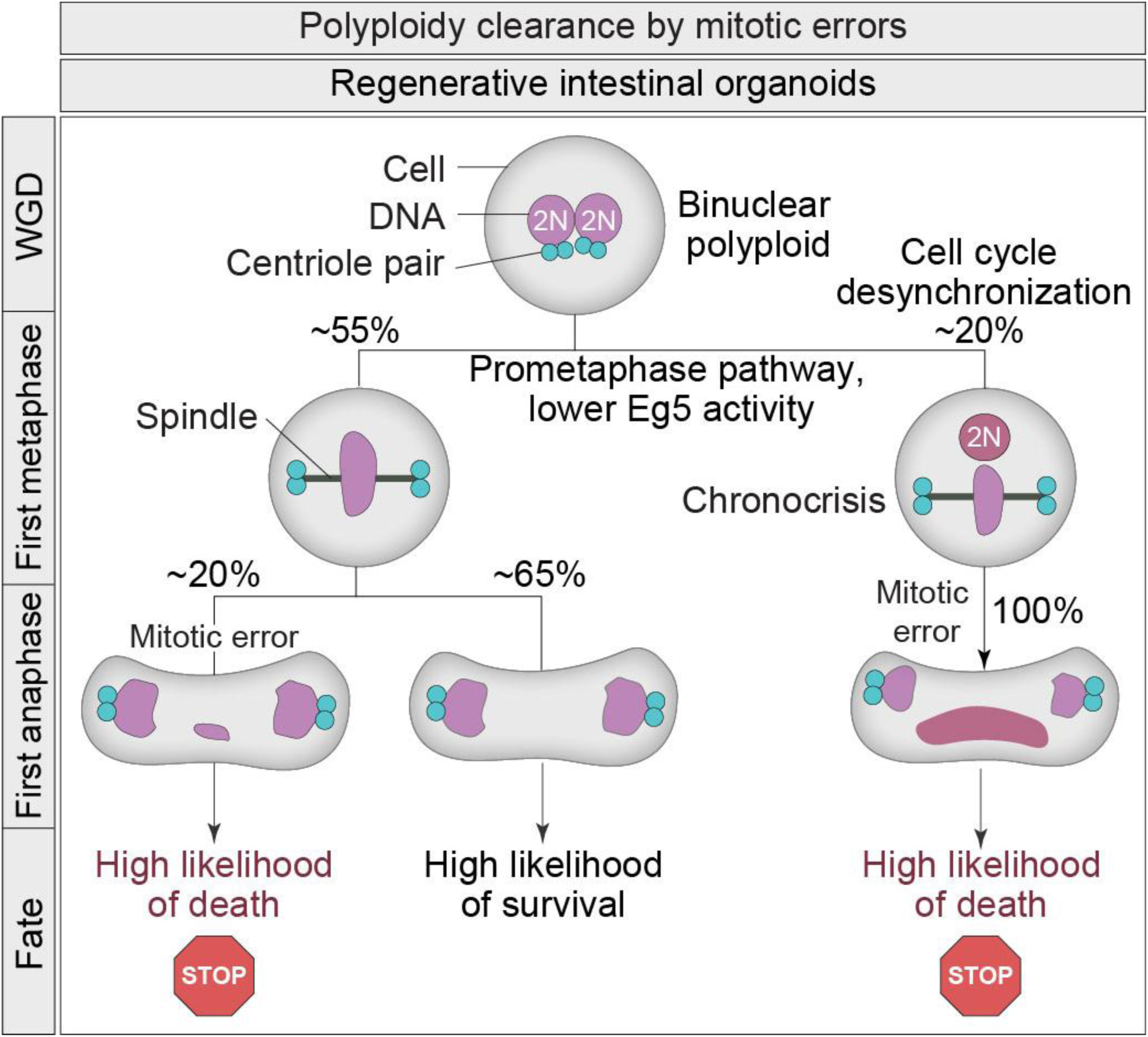
Proposed model for the rapid elimination of proliferative polyploids via mitotic errors in regenerative mSI organoids. All binuclear polyploid mSI cells enter mitosis and in most cases (∼75%) form bipolar spindles, while the remaining divisions involve multipolar spindle configurations (not shown). Among all divisions, ∼55% are bipolar without chronocrisis (left branch), whereas ∼20% are bipolar but display chronocrisis (cell cycle desynchronization of two nuclei, right branch). In bipolar divisions without chronocrisis, cell fate depends primarily on chromosome segregation fidelity: small chromosome missegregation events (20% of bipolar divisions without chronocrisis) lead to a high likelihood of cell death (STOP sign), while error-free divisions (65% of bipolar divisions without chronocrisis) allow binuclear polyploid cells to propagate for at least two generations if no errors occur. Bipolar divisions with chronocrisis typically result in severe mitotic defects and cell death (STOP sign). Multipolar divisions also frequently lead to cell death (not shown). For simplicity, all bipolar divisions are shown with the dominant symmetrical centrosome clustering configuration (2:2), which occurs in 70% of bipolar divisions. No interphase arrest associated with extra centrosomes was observed during early organoid growth and is therefore not shown.

### Centrosome checkpoints, YAP signalling and depolyploidization in regenerative mSI

Recent studies suggest that regenerative mSI tissue may permit transient proliferation of polyploid cells^26^, expanding the view that the liver is the only healthy organ where polyploids routinely cycle^18^. Our results reinforce this idea. Typically, cells with extra centrosomes activate the Hippo pathway^9,68^ or p53 via the PIDDosome^33^, but early mSI organoids show no immediate arrest after polyploidization (Figure 4A) and we even observed dividing octaploids (Figure 1 and 3). As Yes-associated protein 1 (YAP1) signalling blocks Hippo activity^9^, the unperturbed cycling of polyploids is consistent with transient YAP1 activation during the first 72 hours of mSI organoid growth^27,69^ and with its density-dependent deactivation later^69^. Progressive YAP1 deactivation may thus act as a slower-acting checkpoint that limits proliferation of any remaining polyploids with extra centrosomes in larger organoids. Together with fast selection against mis-segregating cells during early growth, this could explain why most polyploids that persist in larger organoids at 96 h have lost their extra centrosomes (Figure 1). In systems like mSI, where clustering is efficient and multipolarity is rare (Figure 1), there may be little selective pressure against extra centrosomes, as opposed to many cell lines^32^, allowing alternative routes of polyploid elimination to dominate. The low incidence of multipolar spindles (Figure 1) also argues against depolyploidization via reduction divisions^21^ as a major clearance route, although rare mononucleated daughters of multipolar divisions can produce viable, dividing progeny (Figure 4), in accordance with recent results on cell lines^70^. While negligible in early organoids, reduction divisions contribute more substantially in contexts where polyploids accumulate in time, such as chronic liver regeneration, where they promote karyotypic diversification^23^.

### Source of mitotic errors in regenerative mSI

We found that the main selective pressure acting on proliferating polyploids in mSI organoids is their frequent chromosome missegregation (Figure 4). In the liver, bipolar polyploids can divide with either high or low fidelity, probably depending on whether regeneration is acute or chronic^20,21,23^. In regenerative-like mSI early organoids, cells that die shortly after polyploidy induction are those that mis-segregated chromosomes in their first division (Figure 4), and chronocrisis events are a major contributor to catastrophic errors (Figure 3). Chronocrisis affects only binucleated cells^15^, so its strongest effect occurs in the first polyploid division, with limited contribution thereafter due to the rarity of new binucleated daughters (Figure 4). Surviving progeny of chronocrisis events may carry extensive DNA damage^14,15^, leading to continued errors. Error rates in uninduced and induced bipolar polyploids are similar (∼33% vs. ∼42%, Figures 3A and 3E) despite differing frequencies of chronocrisis (∼7% vs. ∼25%, Supplementary Fig. 4G and Figure 3E), supporting this view. On the other hand, many non-catastrophic errors (∼17% in the first division) likely arise from merotelic attachments^71^ during pseudo-bipolar divisions^13^, similar to non-tumor cell lines after induced polyploidy (∼10%)^13^. Replication stress after a single polyploid S-phase may also contribute^8^. These differing error types likely explain the variable proliferation outcomes observed after induced polyploidization in mSI organoids (Figure 4), similar to prior study on human colonoids^72^. Consistent with our progeny tracking results, a previous study on human patient-derived colonoids^73^ found mitotic errors are often linked to cell death of progeny. As cell death after mitotic errors was linked to p53 proficiency status in some but not all patients^73^, it will be important to determine if death after mitotic errors during intestinal regeneration is p53-dependent.

Missegregation in mSI organoids does not appear to result from defects in apicobasal polarity (Figure 3D and Supplementary Fig. 4A-B), in contrast to primary mammary spheroids^20^. Intestinal cells are highly polarized^74^ and we found that this occurs very early, as central lumen is formed as soon as the two-cell stage (Supplementary Fig. 4A-B), whereas mammary spheroids polarize later^20^. mSI organoid-derived cells maintain similar error rates in organoids and monolayers (Figure 3C), indicating that errors are cell-intrinsic and linked to excess DNA and centrosomes. Notably, the fraction of cells with extra centrosomes is far lower in immature and mature mammary spheroids (∼6% and ∼4%)^20^, than in early mSI organoids (∼16%, Figure 1), highlighting differences in the mitotic challenges faced by these epithelial systems. Accordingly, it will be interesting to explore if polyploidy arises during acute or chronic regeneration in native intestinal tissue, and how well mitotic fidelity is maintained in this context.

### Mechanisms of rapid clustering of extra centrosomes in regenerative mSI

Tissue-specific features also underlie the rapid clustering of extra centrosomes in mSI compared to non-tumor cell lines^12^. In mSI cells, incomplete centrosome separation at mitotic onset and low Eg5 activity (Figure 2) likely facilitate clustering. Incomplete separation has been associated with faster mitosis and modestly increased segregation errors in cancer cell lines^45^. We propose that this phenomenon also drives efficient clustering in a healthy tissue-mimicking context. Incomplete centrosome separation may promote clustering by keeping centrosomes within the HSET-dependent clustering range and increasing the time they remain near each other^39^. Centrosome–nucleus geometry may also contribute: before mitosis, centrosomes are anchored apically while the nucleus migrates toward them in mSI organoids^75^, potentially reducing chromosome interference with clustering^59^. In agreement, the loss of interphase centrosome anchoring increases spindle multipolarity in Caco-2 cells^65^. Low Eg5 activity directly promotes clustering by weakening outward forces that oppose minus-end–directed motors^41–43^. As in other systems^38,53^, the velocity of clustering of extra centrosomes strongly correlates with mitotic duration (Figure 2). However, polyploid mSI cells divide much faster than reported for polyploid cell lines, consistent with highly efficient and rapid clustering in this system. However, given links between mitotic timing, checkpoint systems and proliferative capacity^61,76^, even a small increase in mitotic duration we observed in polyploids compared to diploids may trigger the “mitotic stopwatch” and restrict proliferation of polyploid progeny on a longer timescale, as recently observed in newly generated polyploids without extra centrosomes^11^. However, the lack of cell cycle prolongation that we observed after polyploidization argues against this mechanism being dominant in limiting proliferation of polyploids during early mSI growth.

### Intestinal inflammation and long-term risk of chronic polyploidization

In conclusion, efficient centrosome clustering in naturally arising mSI polyploids prevents multipolar divisions, but high missegregation rates trigger rapid elimination before slower-acting mechanisms can act. When multipolarity is rare, selection against extra centrosomes is insufficient to drive positive selection for centrosome loss^32^. Instead, polyploidy in mSI appears to arise from a fetal-like regenerative gene expression program^26^, but its low frequency in early mSI organoids suggests it does not confer a strong tumor-protective role, unlike in hepatocytes^22,77^. Instead, gene expression changes in mSI may support regeneration without overwhelming the tissue with cytokinesis-failure–induced polyploids. This is consistent with our observation that substantial early-stage enrichment of polyploidy fails to support efficient organoid maturation (Figure 4). Under chronic regeneration, however, polyploid cells may arise persistently and create new routes for pathological progression. As chronic inflammation is strongly associated with cancer development^78,79^, and polyploid cells are present in ulcerative colitis^80^, we speculate that polyploidy removal by mitotic error surveillance pathway also operates in chronic intestinal inflammation.

## Materials and Methods

### Organoid lines

#### Murine small intestinal organoids

Mouse small intestine (mSI) organoids were obtained from the small intestines of female wild-type C57BL/6 mice as previously described^37^. These organoids were genetically modified by lentiviral transduction to stably express H2B-mNeonGreen using the pLV-EF1a-H2B-Neon-IRES-Puro plasmid^81^, similarly to previous work^82^. The generation of mSI organoids with stable expression of both EGFP-Centrin-2 and Cre-inducible H2B-mCherry was previously described^37^.

#### Human colonic organoids

Normal colon tissue, isolated from a human donor, was provided by the Netherlands Cancer Institute Antoni van Leeuwenhoekziekenhuis Biobank. Tissue was identified as normal based on pathological assessment before establishing human colon (hC) organoids. These organoids were then genetically modified to stably express H2B-mNeonGreen as previously described^82^. Informed consent was acquired from the tissue donor prior to receiving the specimen used in this study.

### Organoid culture

#### Mouse small intestinal organoids

To obtain the intestinal crypts, a freshly dissected small intestine of a mouse was cleaned and cut into small pieces in cold PBS. Pieces of tissue were then gently mixed in cold 20 mM EDTA in PBS for 1h to dissociate the cells and then transferred to a fresh EDTA solution. For six iterations the tissue pieces were incubated in fresh, cold EDTA solution for 5-10 minutes and shortly agitated before passing the resulting suspension through a 70 µm cell strainer. Strained suspension with the best quality and quantity of the crypts was centrifuged at 300 g for 7 min at 4°C. Resulting crypt pellet was resuspended in cold PBS and a portion of the suspension was mixed with culture medium and in a 1:2 ratio with Matrigel (#356255, Corning). This mixture was plated in pre-warmed 24-well plates as 30 µl domes and solidified for 10 min at 37°C and in 5% CO_2_. All mSI organoids were maintained in IntestiCult™ Organoid Growth Medium (Mouse, #06005, STEMCELL Technologies). Passaging of all mSI organoids was performed using mechanical fragmentation by pipetting. To dissociate the cells and establish cystic organoids for experiments, organoids were mechanically fragmented by pipetting and washed in Advanced DMEM/F-12 (Gibco). After incubation of the pellet in TrypLE™ Express (#12604013, Gibco) for 5 min at 37°C, resulting suspension of single cells and small clumps of cells was washed in Advanced DMEM/F-12 and centrifuged at 300 g for 3 min. Resulting pellet was mixed with culture medium and Matrigel in a 1:2 ratio and plated on 35 mm uncoated dishes with 0.17 mm glass thickness (Ibidi) as 2 µl domes and covered with 500 µl of the IntestiCult™ medium after Matrigel solidification at 37°C. Samples were fixed at 48h and 96h post seeding, to enrich for small and large organoids, respectively. Large organoid samples seeded for live-imaging, were prepared in a similar manner, but without the chemical dissociation step, and imaged 24h after seeding.

For the establishment of the intestinal epithelial monolayer culture, dissociated organoids were densely seeded onto a Matrigel layer prepared by coating a well of a CELLview 4-compartment 35-mm glass-bottom dish (Cellview) with 50 µl of a Matrigel diluted in culture medium to a final concentration of 6 µg/µl. The Matrigel layer was allowed to solidify for 30 min at 37°C prior to seeding. In monolayer culture, 10 µM ROCK inhibitor Y-27632 (HY-10583, MedChemExpress) was added to the culture medium.

#### Human colonic organoids

Establishing the culture of human colonic organoids was done as described before^83^. Organoids were maintained in IntestiCult™ Organoid Growth Medium (Human, #06010, STEMCELL Technologies) and passaged by mechanical fragmentation by pipetting. For the establishment of the hC cystic organoids for experiments, the same procedure was used as for mSI cystic organoids.

### Small molecule inhibitors

#### Inducing polyploidy

To induce cytokinesis failure, mSI organoids were incubated for 22h with 1 µM cytochalasin D (HY-N6682, MedChemExpress) 48h or 96h after seeding dissociated organoids. Cells were then washed five times with 2 ml of warm Advanced DMEM/F-12 (Gibco) before adding fresh culture medium and conducting one of the following: 1) fixed after 4h or 2) 48h of incubation at 37°C, 3) transferred to a heated chamber of a Lattice Lightsheet 7 microscopy system for live-imaging or 4) further treated with inhibitors.

#### Motor protein and microtubule polymerization inhibitors

Inhibitors of motor proteins were used at following concentrations and incubation times: monastrol (HY-101071A, MedChemExpress), 30 µM for 3h; CW069 (HY-15857, MedChemExpress), 250 µM for 30 min or 400 µM for 1h. Nocodazole (HY-13520, MedChemExpress) was used at 3 µM concentration for 2h to fully depolymerize microtubules. In nocodazole washout experiments, after the 2h incubation, cells were washed as described above and fixed after 45 or 60 minutes of incubation with fresh culture medium at 37°C.

### Immunofluorescence

Mouse SI and hC organoids were fixed with warm 4% paraformaldehyde at 37°C for 1h and monolayers for 20 min. A portion of mSI cells in Supplementary Figure 3A and 3D (“anaphase and telophase cells”) and mSI organoids stained for integrin β4 were fixed with ice-cold methanol for 1h at 4°C. Permeabilization, blocking and working buffer were prepared as described previously^55^. Primary antibodies were diluted as follows: rat anti-α-tubulin (1:100, MA1-80017, Invitrogen), rabbit anti-centrin-3 (1:500, ab228690, Abcam), rabbit anti-ZO-1 (1:100, #61-7300, Invitrogen), rabbit anti-integrin β4 (1:50, ab182120, Abcam), mouse anti-E-cadherin (1:500, ab231303, Abcam). Primary antibodies were incubated with cells overnight at 4°C. Secondary antibodies, all diluted to 1:500, were as follows: donkey anti-rat Alexa Fluor 594 (ab150156, Abcam), donkey anti-rabbit Alexa Fluor 488 (ab150061, Abcam), donkey anti-rabbit Alexa Fluor 594 (ab150064, Abcam), donkey anti-mouse Alexa Fluor 488 (ab150105, Abcam). Secondary antibodies were incubated with cells overnight at 4°C in combination with SiR-actin (1 µM, Spirochrome) for F-actin staining. DNA was stained with DAPI (1 µg/mL, Sigma-Aldrich). In the organoid formation assay experiment, samples were fixed with 4% paraformaldehyde, stained overnight with 1 µM SiR-actin (Spirochrome) and shortly with DAPI (1 µg/mL).

### Microscopy

#### Fixed-cell imaging

Single cells and organoids stained for polarity factors were imaged with the Expert Line easy3D STED microscope system (Abberior Instruments) with the UPLSAPO 100x/1.4 NA oil objective (Olympus) and avalanche photodiode (APD) detector. Laser lines of 405 nm, 488 nm, 561 nm, or 647 nm were used for excitation. Images were acquired using the Imspector 16.3 software. The xy pixel size was set at 100 nm and z-step size at 500 nm for imaging an individual cell’s field of view and 200nm with a z-step size of 2 µm for images of the whole organoids.

Mitotic cells within untreated mSI organoids were also initially imaged with the Expert Line easy3D STED microscope system, as described above. After establishing that no information is lost when xy pixel size is decreased, the LSM 800 confocal laser scanning microscope system (Carl Zeiss) was used to increase the speed of imaging by using the multiposition imaging mode. Images were acquired using the Plan-Apochromat 63x/1.4 NA oil DIC M27 objective (Carl Zeiss) and GaAsP detectors. Laser lines of 405 nm, 561 nm, and 640 nm were used for excitation. Image acquisition was done with ZEN 3.7 (blue edition) software (Carl Zeiss). The xy pixel size was set to 200 nm and z-step size to 500 nm, while the z-stack size was ranging between 60-80 μm, depending on the average organoid size in each experiment. The same microscope system was used for all fixed-cell experiments, including human colonic organoids stainings.

Organoid formation assay samples were imaged using the Andor Dragonfly spinning disk confocal microscopy system (Andor Technology) with the Plan Fluor 40x/1.3 NA oil objective (Nikon), with excitation laser lines of 488 nm and 637 nm and Sona sCMOS camera (Andor Technology). Fusion 2.6.0 software (Andor Technology) was used for image acquisition with the Montage mode enabled and set to Edge mode with 10% overlap. The x and y positions in the Montage mode were set to encompass the whole Matrigel dome. The z-step size was set to 2 μm and the z-stack size to 100 μm.

#### Live-cell imaging

Mouse SI organoids and large organoids expressing EGFP-Centrin-2 and H2B-mCherry were imaged using the Andor Dragonfly spinning disk confocal microscopy system (Andor Technology) with the Plan Fluor 40x/1.3 NA oil objective (Nikon). Laser lines of 488 nm and 561 nm were used for excitation. Simultaneous acquisition of the two channels was accomplished using a dual-camera system consisting of Sona sCMOS camera and iXon Ultra 888 EMCCD camera (Andor Technology) with the use of 565 nm LP image splitter. Fusion 2.6.0 software (Andor Technology) was used for image acquisition. The z-step size was set to 1 μm and time interval to 1 minute. For the duration of imaging, cells were kept at 37°C and 5% CO_2_ within the heating chamber (Okolab).

#### Long-term live-cell imaging

For long-term imaging of control and polyploidy-induced mouse SI organoids expressing mNeonGreen-H2B, the Lattice Lightsheet 7 system (Carl Zeiss) was used, with an 13.3x/0.4 NA illumination objective lens (at a 30° angle to cover the glass), a static phase element and a 44.83x/1.0 NA detection objective lens (at a 60° angle to cover the glass) with an Alvarez manipulator. A laser line of 488 nm was used for excitation with laser power set to 0.2%. The detection module consisted of a Hamamatsu ORCA-Fusion sCMOS camera with exposure time set to 10 ms. Images were acquired using ZEN 3.7 (Carl Zeiss). The light sheet’s dimensions were set to 100 µm x 1400 µm or 100 µm x 1800 µm. The width of the imaging area in the x dimension was set to a range of 100-150 mm, with the x interval of 0.4 µm. The time interval was set to 2 minutes. The total imaging duration was 48 h, or shorter if the acquisition was interrupted by an air bubble, formed during automatic immersion of water. During imaging, cells were kept at 37 °C and at 5% CO_2_ in a heated chamber (Carl Zeiss). Before image analysis, raw images were deskewed using ZEN 3.7 software with the “Linear Interpolation” and “Cover Glass Transformation” settings.

### Image and data analysis

Image analysis was performed in Fiji/ImageJ (National Institutes of Health), unless otherwise specified. Raw images of both fixed and live-imaged cells were used for quantifications. Quantitative analysis of the data was performed using custom-made scripts in MATLAB 9.10.0 (MathWorks) or functions in Excel (Microsoft). In order to determine the mitotic phase and prometaphase morphology in tilted cells, Imaris Viewer 9.8.1. was used (Andor Technology).

#### Organoid size

Organoid size was determined by measuring the largest diameter of the organoid in the maximum intensity projection of all imaged channels. Organoid size categories were defined by following diameters: small organoid, <60 µm; middle-sized organoid, 60-90 µm; large organoid, >90 µm. Cells coming from organoids with less than a half of the volume imaged in the z dimension were excluded, unless their size already exceeded 90 µm (or 70 µm in experiments with combined untreated anaphases and telophases).

#### Organoid morphology

An organoid was categorized as “columnar”^56–58^ if all the interphase cells in its middle z-plane had nuclei at least [0.5 x nucleus length] away from the apical actin border. Cells that began their migration towards the apical side or were returning to the basal side, as evidenced from the detachment from the basal actin border, were excluded from examination. For organoids larger than 70 µm a single-cell deviation from the established rule was tolerated due to the large number of examined cells. Large organoids that could not fit in the field of view were included in the analysis only if the ∼70-80% of the organoid cells were in the middle z-plane field. All the organoids that failed to meet these criteria, including cuboidal^56^ and squamous^56,58^ organoids, were assigned as “non-columnar”.

#### Centrosome number and spindle polarity

In fixed mSI cells, centrosome number was measured as the number of EGFP-Centrin-2 foci (pairs or overlapped) that co-localized with the signals of the tubulin poles and were contained within the cell bounds (determined by the F-actin staining). Clusters of three centrin foci were considered to represent a single centrosome (i.e. a centriole pair with an associated centriolar satellite). In live-imaged mSI cells with centrin labelling, centrosome number was determined solely as the number of EGFP-Centrin-2 foci that showed directed movement during mitosis, starting from the apical border of the organoid (evident by the interphase centrin foci lining) and ending up behind the separating chromosome groups in anaphase.

In fixed cells with two centrosomes, spindles were assessed as bipolar when they contained only two tubulin poles, normally co-localizing with centrin foci in untreated cells. In cells with more than two centrosomes, spindle polarity was assigned as bipolar when all the neighbouring centrin foci within the two centrosome clusters were separated by less than 6 µm and as multipolar when at least one neighbouring distance exceeded 6 µm. In a subset of cells with more than two centrosomes, a single or multiple centrin foci co-localized with weak tubulin signals with or without adjacent chromosomes or chromosome groups. In these cases, when the weak pole was further than 6 µm away from a regular spindle pole, but attracted no chromosomes, the spindle was considered multipolar with an inactivated pole and separated from the previously multipolar spindle category. In fixed anaphase and telophase cells lacking EGFP-Centrin expression, spindle polarity was determined based on the number of chromatin groups; where possible, pole identities were confirmed using immunostained centrin-3 foci, and centrosome numbers were not determined due to high non-specific staining. In the live images of mSI cells expressing either H2B-mNeonGreen or H2B-mCherry, spindle polarity was determined by the number of emerging chromatin groups in anaphase or telophase, respectively. Divisions in which the number of emerging chromatin groups changed from three at anaphase onset to two in later anaphase were separated from “multipolar” cell category to “multipolar-bipolar” category. Divisions in which a third chromatin group remained passive during anaphase were not assessed as multipolar due to a potential chronocrisis event. In cells expressing H2B-mCherry, additional EGFP-Centrin signal was used to confirm that bipolar cells had clustered centrosomes closer than 6 µm before anaphase onset.

#### Segregation errors

Separation of anaphase from telophase cells^84^ was based on the following criteria: opened cleavage furrow (determined by the F-actin staining); chromosome groups appearing condensed and not yet finely rounded; tubulin spindle poles still apparent; absence of the tubulin midbody.

All the anaphase cells were considered to have a segregation error^85,86^ if at least one of the following events occurred within the cell bounds (determined by the F-actin staining): any chromatin in between and not in contact with the segregating groups; any chromatin connecting the segregating groups (unless in very early anaphase), including when there was an interruption in the stretched chromatin connecting the groups; any chromatin on the pole side of the chromosome group, that is visibly separated from the chromosome group (i.e. unaligned polar chromosome^87^). When telophase cells were assessed for segregation errors, the same criteria applied, with the inclusion of cells in which the chromatin strands protruded out of the finely rounded, decondensing chromosome groups.

In live-imaged cells expressing H2B-mNeonGreen, a cell was assessed as having a segregation error if in at least one anaphase frame it had one of the following: any chromatin lagging in between and not directly contacting the segregating chromosome groups; any chromatin connecting the separating chromosome groups, unless in very early anaphase; any stretched chromatin arms that are protruding out of the segregating chromosome groups and reach the midzone between those groups or cause the tilting of the those groups in subsequent frames; a chromatin group separated from the segregating chromosome groups that remains passive during anaphase onset and can prevent anaphase progression. The 3D View option in ZEN software was used to determine chromosome segregation fidelity status of all cells tilted in the z-plane and to confirm the ambiguous status of non-tilted cells. Fidelity status was assessed for all the cells that had a clear signal of chromatin in the first division after polyploidization.

#### Cells in chronocrisis

An anaphase cell was defined as being in chronocrisis^14,15^ if it had either a chromatin group without an adjacent spindle pole, or scattered chromatin mass on the side of the spindle, sometimes partially incorporated into the spindle. The level of DNA condensation of the pole-less chromatin group often differed from the groups on the spindle, but this difference was not always clear and therefore not a defining criterion.

#### Cell and spindle size parameters

Spindle length in metaphase cells was measured as the three-dimensional distance between the centrin signals of the two opposing centrosomes or the midpoints of centrosome clusters, using Pythagorean theorem. The z-dimension distance was multiplied by a factor of 0.85 to account for the refractive index mismatch between aqueous samples and immersion oil^88^.

Metaphase plate length was measured on the maximum intensity projection (MIP) of the z-stack encompassing the whole plate, as the distance between the outermost chromosome edges, perpendicular to the centrosome axis. When the overlapping DNA signal of the neighbouring cells prevented measurement in the MIP, measurement was done by manually identifying the two z-planes containing the outermost chromosome edges and measuring the distance between xy locations of the edges.

Cell circumference of metaphase and anaphase cells was measured using the segmented line tool to outline the cortical signal of F-actin in a single z-plane that was roughly equidistant from the z-planes of opposing centrosomes or centrosome clusters. In cells with multipolar spindles, measurement was done in the center of the cell, determined as the midpoint between either the outermost F-actin cortical signals or spindle signals, in the z-dimension. Anaphase cells that had centrosomes spanning over more than 20 z-planes were considered to have a large tilt and were thus excluded from this analysis due to measurements reflecting the circumference in the transversal plane, instead of the longitudinal plane of the naturally elongated anaphase cell.

#### DNA volume

Segmentation of imaged DNA channels was conducted in Python 3.12.0 (Python Software Foundation) using the public pyclesperanto 0.17.0 GPU-accelerated image processing library (https://github.com/clEsperanto/pyclesperanto). The library implements the Voronoi-Otsu labeling algorithm which applies Gaussian smoothing (spot sigma = 1-15 pixels; outline sigma = 1 pixel), Otsu thresholding, and Voronoi-based region expansion to generate labeled objects. Each segmentation output was manually validated using the 3D Manager plugin in Fiji/ImageJ and volume of the region of interest was measured. To detect polyploid cells with two centrosomes, a minimum threshold value for the increased DNA volume was set as the mean + SD value of cells with two centrosomes (dashed line in Figure 1C and shaded area in Supplementary Fig. 3C). To detect polyploid cells in bipolar anaphases and telophases with segregation errors, a minimum threshold value for the increased DNA volume was set as the mean - SD value of multipolar cells (shaded area in Supplementary Fig. 3D).

#### Spindle polarity and centrosome distances in drug treatments

Spindle polarity after nocodazole washout and in CW069-treated cells was assessed in different mitotic phases, as described before for untreated cells. In both treatments, spindle poles without co-localized centrin sometimes appeared and they were accounted for in the assessment of polarity with the same criteria, if four centrosomes were present in the cell. In the nocodazole washout experiment, pre-anaphase cells were defined as all mitotic cells ranging from those that formed early mitotic spindles to cells in metaphase.

Centrosome distances in cells with two centrosomes arrested in nocodazole or monastrol were calculated as was described for spindle length in metaphase. In cells with four centrosomes, centrosome distance was calculated similarly, by measuring the centrosome coordinates and calculating unique pairwise distances using MATLAB. Centrosome identity confirmation via tubulin, described before, was not possible in nocodazole-treated cells due to the full microtubule depolymerization. Monastrol- and nocodazole-treated cells were roughly categorized as small or large, if their cell size was smaller or larger than 56 µm, based on the cutoff values of metaphase cell size in cells with two and four centrosomes (Supplementary Fig 1D).

#### Centrosome position tracking

Centrosome positions were manually tracked using the multi-point tool in Fiji/ImageJ to mark the centrin foci in each frame. Centrosome identity was confirmed as described above for centrosome number analysis in live images. The tracks were then categorized based on the centrosome counts, and DNA volume: cells with nuclei volumes smaller or larger than mean + SD value of cells with two centrosomes were considered as “normal” or “large”, respectively, to separate the polyploids with potential centrosome loss. DNA frames chosen for segmentation (*see “DNA volume”*) corresponded to interphase nuclei five minutes before NEB. Quantitative analysis of centrosome dynamics, including NEB distance, metaphase distance and maximum distance was performed using custom-made MATLAB scripts.

#### Progeny tracking

Cells in live-imaged mSI expressing mNeonGreen-H2B were tracked only if they entered mitosis after imaging was initiated. When polyploidy was induced, only cells that entered their first mitosis as a binuclear were chosen for analysis. ZEN 3.7 software was used to manually follow an individual cell for at least one generation (until “daughter outcome”). Other than division, possible progeny outcomes also included death and the unknown outcome. The former was defined as the event in which the cell’s nuclear architecture suddenly starts changing and the cell migrates into the organoid’s lumen or is extruded through the basal surface. Cells that died in mitosis were very rare and thus excluded from analysis. Cells that died >43h after their mother division were excluded from the death outcome, due to possible phototoxicity effects^73^. Unknown outcome was any event in which the progeny outcomes were impossible to determine due to blurry or missing organoid regions or imaging end.

Cells that did not divide for >30h after their mother division were separated from the unknown category and termed “non-proliferative” in daughter outcomes. Cells that were followed for <8h after their mother division were excluded from this analysis. In short videos, in which it was impossible to determine any progeny outcome, only parameters of the first cell division were analysed (*see* “*Mitotic and cell cycle time”, “Segregation errors” and “Centrosome number and spindle polarity”)*.

#### Mitotic and cell cycle time

In live images of mSI cells, mitotic time was defined as the time elapsed from NEB to anaphase onset. In cells expressing mNeonGreen-H2B, the frame of NEB was defined as the last frame in which chromosomes are contained in a shape of the nucleus, and anaphase onset was detected by the sudden separation of chromosomes within the metaphase plate. In cells expressing H2B-mCherry and EGFP-Centrin-2 the frame of NEB was defined as the first frame in which the previously clear DNA signal of a nucleus disperses. Anaphase onset was determined from the dynamics of centrosome separation (*see “Centrosome position tracking”*) and defined as the last frame with a stable metaphase centrosome distance, after which a continued increase in centrosome distance ensued. Cells that transitioned from spindle multipolarity to bipolarity in anaphase were rare and thus excluded from the comparisons of mitotic timing based on polarity.

In mSI organoids expressing mNeonGreen-H2B, the cell cycle time was defined as the time elapsed between the NEB of a mother cell and the NEB of its daughter cell.

#### Organoid formation assay

In organoid formation assays, organoid types were quantified from maximum intensity projections of the central 40 µm of the imaged Matrigel dome. Organoids were classified into four categories based on morphology, using F-actin and DAPI staining: budding, cystic, irregular, and dead. Budding organoids were defined as structures exhibiting at least one pronounced epithelial protrusion or as elongated structures with a visible lateral constriction, both indicative of a developing or established crypt-like region. Cystic organoids were defined as structures containing a clearly defined central or dominant lumen, enclosed by a circular or semi-circular epithelial layer. Irregular organoids were defined as structures lacking a single dominant lumen, including those with multiple lumina or with mono- or multilayered cellular patches forming at the dome edges. Small structures consisting of fewer than ∼3 cells were excluded from the analysis, as early-stage organoids do not display pronounced lumina. Dead organoids were defined as clusters of cells exhibiting apoptotic nuclei, accompanied by co-localizing F-actin signal appearing diffuse, punctate, or consistent with membrane blebbing or cellular swelling. Dead cell clusters in direct contact with larger, viable organoids were excluded, as these may represent extruded material from ruptured cystic structures.

### Image processing, data representation and statistical analysis

All the figures were assembled in Adobe Illustrator (Adobe Systems). The schematic in Figure 1F was created using BioRender. The minimum and maximum intensities are not equally adjusted throughout the images, since no comparative measurements were made, but were set to provide the best signal of the imaged phenomena. When indicated, the smoothing of the images was performed using the “Gaussian blur” function in ImageJ (s = 0.5-1.0). Color-coded maximum intensity projections of the z-stacks were made using the “Temporal color code” tool in Fiji/ImageJ by applying “Rainbow RGB” lookup-table (LUT).

All the data plots were assembled in either MATLAB or Excel. For the generation of univariate scatter plots, the open MATLAB extension “UnivarScatter” was used (https://github.com/manulera/UnivarScatter).

When plotting centrosome distance over time, the mean lines with shaded areas representing standard error of the mean were plotted to encompass all of the data points.

The Shapiro-Wilk test was used to determine normality of the sample data, followed by the T-test for data that followed a normal distribution or Mann-Whitney U test otherwise. When comparing data from multiple classes, the one-way analysis of variance (ANOVA) test followed by pairwise Tukey Honest Significant Difference (HSD) test was used for data that followed normal distribution and Kruskal-Wallis test with post hoc Dunn’s test was used otherwise. When comparing frequencies, a two-tailed two-proportion Z-test was used, with overall differences assessed by a chi-square test of independence. In all analyses, the result was considered statistically significant when the p-value was equal to or less than 0.05. Statistical analysis was performed as noted in the respective Figure captions.

## Acknowledgements

We thank the Laboratory for Neurodegenerative Disease Research led by Silva Katušić at the Ruđer Bošković Institute and Ana Petelinec for assistance with isolation of the mouse small intestine; Dragomira Majhen from Laboratory for Cell Biology and Signalling at the Ruđer Bošković Institute for providing conditions for lentiviral work; Lovro Gudlin for adapting the code for segmentation analysis; Alessandro Corsini for technical advice; Eloise van Kwawegen for help with organoid culture; Ivana Šarić for the illustrations and assembling the figures; the University of Zagreb University Computing Centre – SRCE for providing imaging data storage; and members of the Tolić and Kops labs for their constructive comments on the study. This work was funded by the European Research Council (ERC-SyG 855158, I.M.T. and G.J.P.L.K.); HRZZ (IP-2024-05-5336, I.M.T.); Swiss-Croatian Bilateral Project (IPCH-2022-10-9344, I.M.T.); projects co-financed by the Croatian Government and the European Union through the European Regional Development Fund - the Competitiveness and Cohesion Operational Programme IPSted (KK.01.1.1.04.0057, I.M.T.) and QuantiXLie Center of Excellence (PK.1.1.10.0004, I.M.T.). M.T. is a Cancer Prevention and Research Institute of Texas (CPRIT) Scholar in Cancer Research, supported by CPRIT grant RR240079.

## Author contributions

I.D. and K.V. designed the experiments, which were performed and analyzed by I.D. Human-derived organoids provided by B.C. and G.J.P.L.K. were established and genetically modified by T.V.R. G.J.P.L.K. and M.T. contributed to the data interpretation. I.M.T. supervised the project. I.D., K.V., and I.M.T. wrote the manuscript. All authors edited and approved the manuscript.

## Data and code availability

Source data are provided with this paper. All other data supporting the findings of this study and codes that were used to analyse and plot the data are available from the corresponding author on reasonable request.

---

**Supplementary Figure 1.**
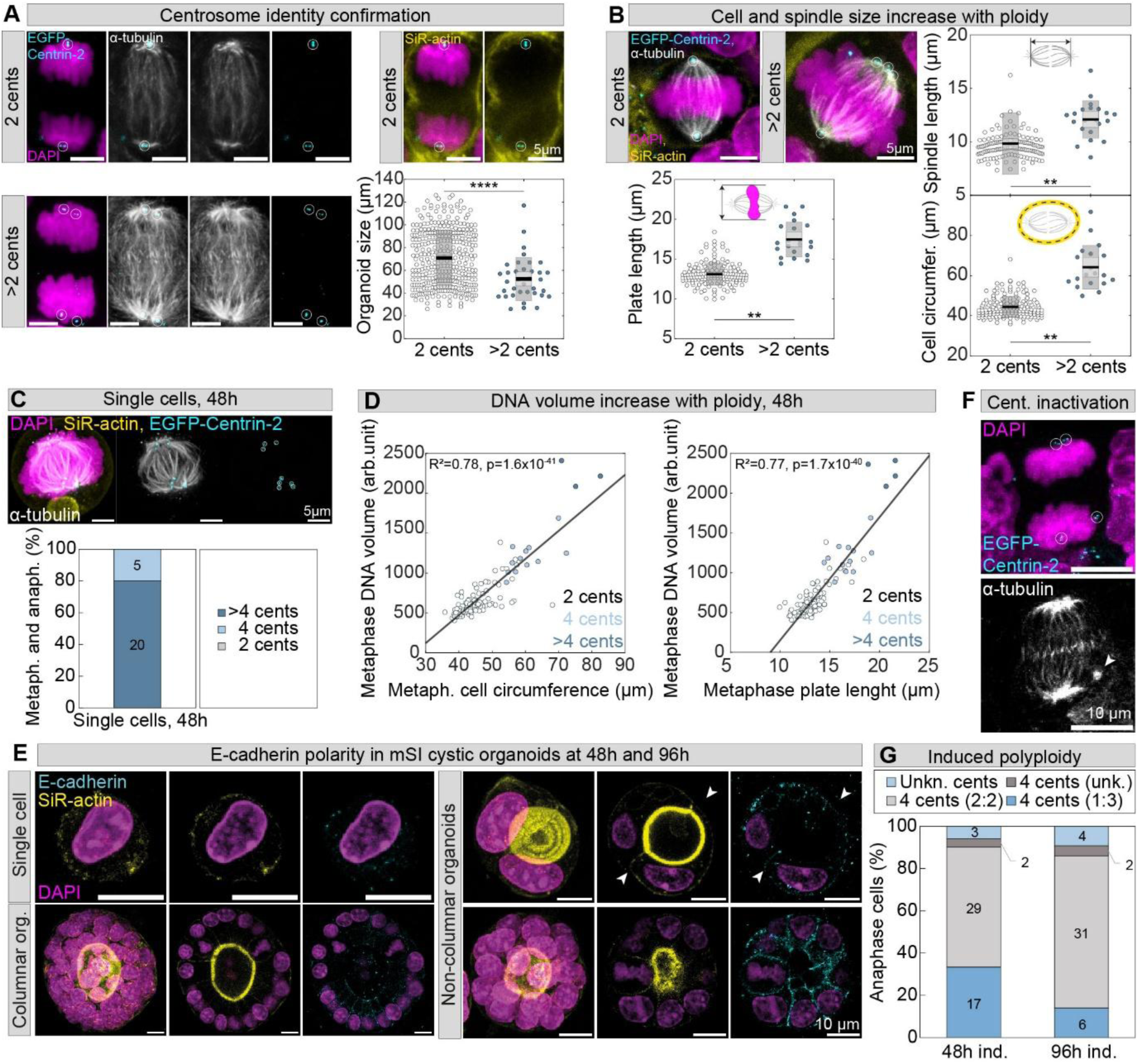
Spindle and metaphase plate dimensions increase with ploidy in mSi organoids with preserved E-cadherin basolateral localization. **(A)** Fixed images of mSI anaphase cells stably expressing EGFP-Centrin-2 (cyan), immunostained for α-tubulin (grey) and stained for F-actin with SiR-actin (yellow) and for DNA with DAPI (magenta). The univariate scatter plot shows the size of each organoid that contained anaphase cells with two or more than two centrosomes. N = (342, 36) cells from (> 300, > 30 organoids) and ≥ 3 experiments. **(B)** Fixed images of mSI metaphase cells with two and more than two centrosomes. The univariate scatter plots show metaphase plate length (N = 152, 21 cells), spindle length (N = 152, 21 cells), and cell circumference for metaphase cells (N = 152, 19 cells) with two and more than two centrosomes. Data from (> 100, > 10 organoids) and ≥ 3 experiments. **(C)** Fixed images of a mSI single cell in metaphase. Graph shows the frequencies of single cells with different number of centrosomes in metaphase and anaphase at 48h post seeding dissociated organoids. N = 25 cells and ≥ 3 experiments. **(D)** Correlation between DNA volume and cell circumference (left) or plate length (right) across metaphase cells with different centrosome numbers. Black lines show linear regression. N = 122 cells from >100 organoids and ≥ 3 experiments. **(E)** Images show fixed organoids and a single cell immunostained for basolateral marker E-cadherin (cyan), stained with SiR-actin (yellow) and DAPI (magenta). The second and third column show single z-planes. White arrows point to cell borders, determined by the F-actin staining. **(F)** Images show an example of a fixed anaphase cell after induction of polyploidy that has a distant inactivated spindle pole (arrowhead). **(G)** The bar plot compares the frequencies of symmetric (2:2) and asymmetric (1:3) clustering configuration of centrosomes in bipolar mSI anaphase cells with four centrosomes. Frequencies are shown for polyploidy induction at both 48h and 96h post seeding. N = (51, 43) cells from > 40 organoids and ≥ 3 experiments. All images, unless otherwise stated, are maximum intensity projections. White circles mark centriole pairs for which centrosome identity was confirmed using α-tubulin staining. In univariate plots, colored points represent individual cells; black lines show the mean, with light and dark grey areas marking 95% confidence intervals for the mean and standard deviation, respectively. Statistics: Z-test for two population proportions when comparing frequencies in (G); Mann-Whitney U test in univariate plots in (A) and (B) and two-tailed t-test in (D). Symbols: **, P ≤ 0.01; ****, P ≤ 0.0001. Cents or cent., centrosome(s); circumfer., circumference; metaph., metaphase; anaph., anaphase; arb., arbitrary; org., organoid; ind., induced; unk. or unkn., unknown.

**Supplementary Figure 2.**
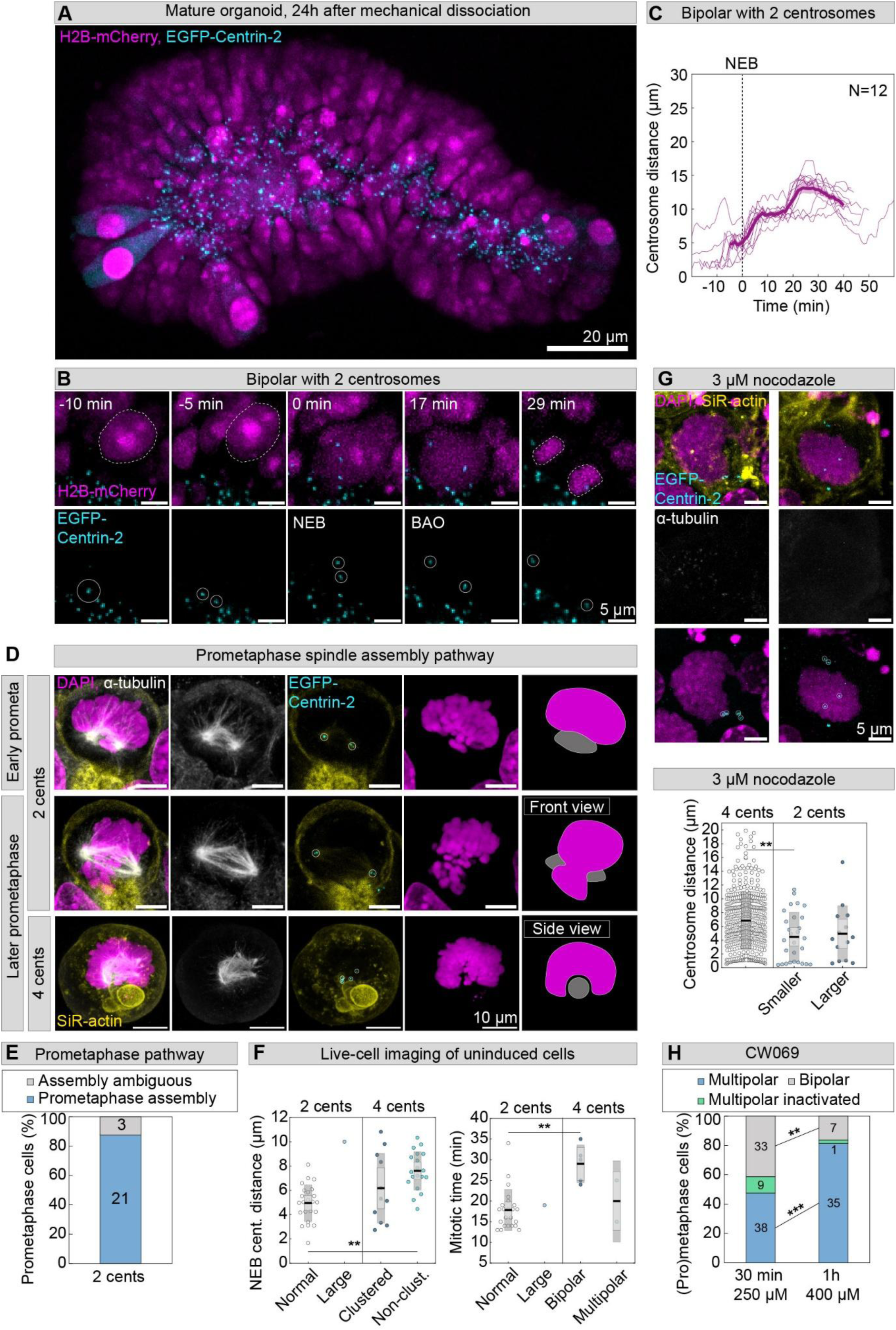
Mitotic cells in mSI organoids assemble spindles via a prometaphase pathway. **(A)** A live-imaged mature mSI organoid, 24h after seeding mechanically fragmented organoids with cells stably expressing EGFP-Centrin-2 (cyan) and with leaky expression of H2B-mCherry (magenta). **(B)** Representative time-lapse images of a bipolar division in a mSI cell with two centrosomes. The cell nucleus and chromatin groups in telophase are outlined with white dashed lines for clarity. **(C)** Plot shows centrosome distances in time (purple thin lines) for all cells with two centrosomes within the mature organoids. N = 12 cells from 2 organoids and 1 experiment. Time zero marks mitotic onset (dashed line); thick lines are mean values; shaded areas are SEM values. **(D)** Fixed images of mSI early and later prometaphase cells with two (first and middle row) or four centrosomes (bottom row) stably expressing EGFP-Centrin-2 (cyan), immunostained for α-tubulin (grey) and stained for F-actin with SiR-actin (yellow) and for DNA with DAPI (magenta). Cells are showing characteristics of the prometaphase spindle assembly pathway, primarily the presence of prometaphase rosettes (DAPI channel, magenta). The last column shows a simplified schematic representation of mitotic spindle (grey) and chromosome (magenta) from the images on the left. **(E)** The bar plot shows the fraction of prometaphase cells with two centrosomes undergoing a prometaphase spindle assembly pathway. N = 25 cells from > 20 organoids and ≥ 3 experiments. **(F)** The univariate scatter plot of centrosome distances at NEB (left) and mitotic time (right) across live-imaged cells from Figure (2A-B) containing two or four centrosomes grouped by nuclear size (“normal”, “large”) and centrosome clustering outcome (“clustered”, “non-clust.”) or spindle polarity in anaphase (“bipolar”, “multipolar”), respectively. N = (23, 1, 5, 2) cells from (> 10, 1, > 3, 2) organoids and ≥ 3 experiments per centrosome group (“2 cents” and “4 cents”). **(G)** Fixed images of mSI anaphase cells treated with 3 µM nocodazole for 2h. The univariate scatter plot shows centrosome distances in cells with four centrosomes treated with 3 µM nocodazole. Six unique distances are plotted for each cell with four centrosomes. Distances in cells with two centrosomes are shown for comparison, with large cells separated, due to possible polyploidy. N = (74, 28, 14) cells from > 50 organoids and ≥ 3 experiments in total. **(H)** The bar plots show the frequency of mitotic spindle polarity groups in fixed mSI anaphase cells treated with 250 µM and 400 µM HSET inhibitor CW069 for indicated durations. In (G-H), prior to treatment, polyploidy was induced using cytochalasin D at 48h post seeding dissociated organoids and only cells with four centrosomes were selected for analysis in (H). N = (80, 43) cells from (> 70, > 30) organoids and ≥ 3 experiments. All images are maximum intensity projections. White circles mark centriole pairs for which centrosome identity was confirmed using α-tubulin staining or by observing centriole pair trajectories during live-cell imaging (see Methods). In (G) centrosome identity is assumed based solely on the centrin signal and cell boundaries. In univariate plots, colored points represent individual cells; black lines show the mean, with light and dark grey areas marking 95% confidence intervals for the mean and standard deviation, respectively. Statistics: Z-test for two population proportions when comparing frequencies in (H); in (F) ANOVA with post hoc Tukey’s HSD test; in (G) Kruskal-Wallis test with post hoc Dunn’s test. Symbols: **, P ≤ 0.01; ***, P ≤ 0.001. NEB, nuclear envelope breakdown; BAO, before anaphase onset; cents, centrosomes; Non-clust., non-clustered.

**Supplementary Figure 3.**
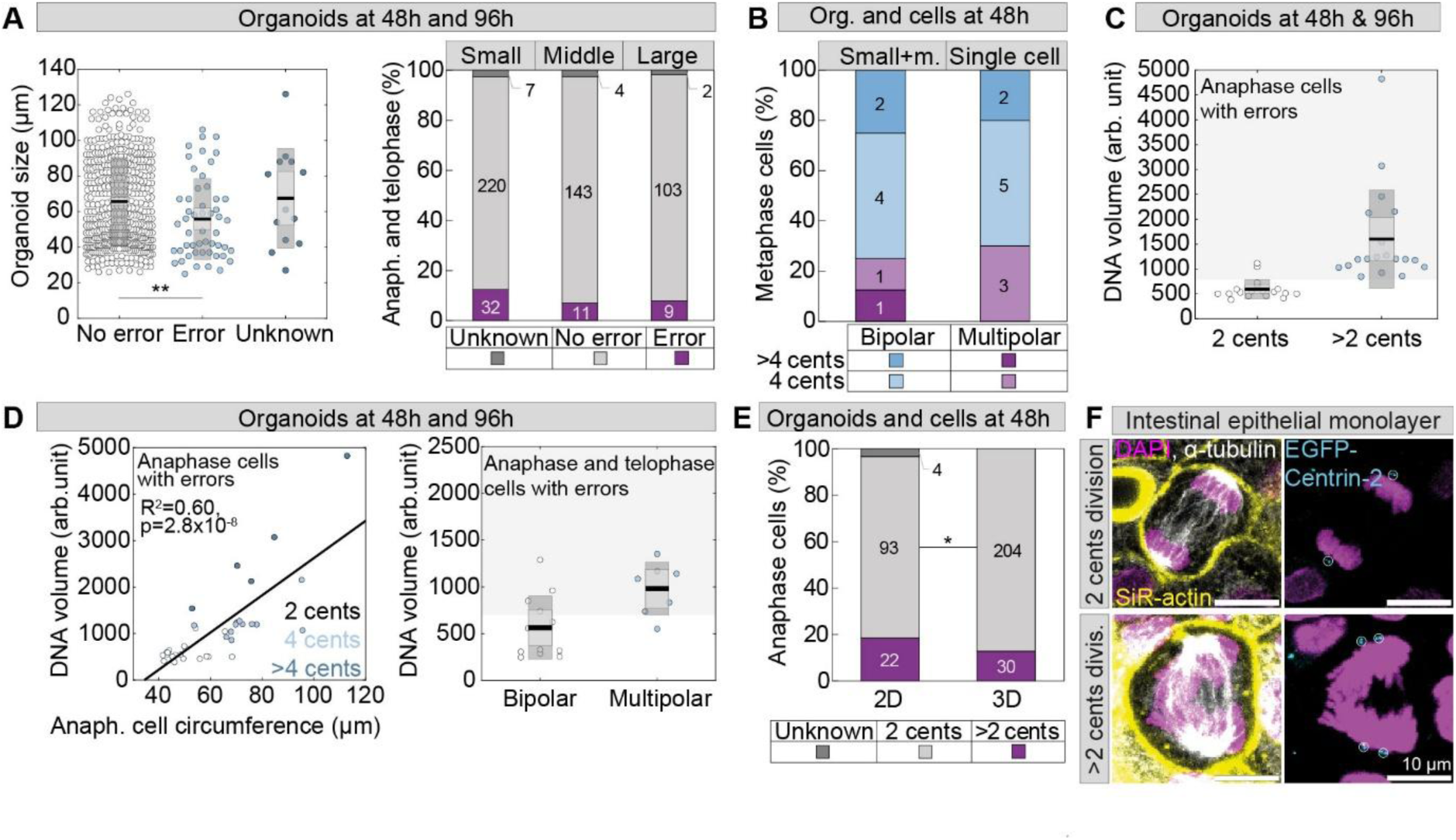
Polyploid cells during early regeneration exhibit anaphase errors independently of organoid architecture formation. **(A)** The univariate scatter plot shows the size of each organoid that contained anaphase cells, separated by chromosome segregation outcomes. N = (453, 50) cells from (> 400, > 40) organoids and ≥ 3 experiments per group (left). The bar plot shows the frequency of chromosome segregation errors of mSI anaphase and telophase cells in organoids of different sizes. Numbers of analysed cells are indicated on the data labels. Data from > 100 organoids and ≥ 3 experiments in each group (right). **(B)** The bar plot shows frequencies of each spindle polarity type in mSI metaphase cells with more than two centrosomes, in organoids and single cells. N = 8 cells from > 5 organoids and 10 single cells, both from ≥ 3 experiments. **(C)** The univariate scatter plot shows the DNA volume of anaphase cells that had a chromosome segregation error, separated by centrosome numbers. Shaded area (grey) marks the values of DNA volume that are above the mean + SD of cells with two centrosomes, indicating polyploidy (see Methods). N = (19, 17) cells from > 15 organoids and ≥ 3 experiments in each group. **(D)** Correlation between DNA volume and cell circumference (left) across anaphase cells with different centrosome numbers. Black line shows linear regression. N = 35 cells from >30 organoids and ≥ 3 experiments. The univariate scatter plot shows the DNA volume of anaphase and telophase cells that had a chromosome segregation error, separated by spindle polarity (right). Shaded area (grey) marks the values of DNA volume that are above the mean - SD of multipolar cells, indicating polyploidy (see Methods). N = (12, 6) cells from > 5 organoids and ≥ 3 experiments in each group. **(E)** The bar plot shows the frequencies of mSI anaphase cells with two or more than two centrosomes, depending on whether they divide in a monolayer (“2D”) or in organoids (“3D”), at the same timepoint after seeding dissociated organoids (48h). N = (119, 234) cells from > 100 organoids and ≥ 3 experiments in each group. **(F)** Fixed images of mSI anaphase cells stably expressing EGFP-Centrin-2 (cyan), immunostained for α-tubulin (grey) and stained for F-actin with SiR-actin (yellow) and for DNA with DAPI (magenta), dividing within a monolayer with two or more than two centrosomes. Images are maximum intensity projections. White circles mark centriole pairs for which centrosome identity was confirmed using α-tubulin staining. In univariate plots, colored points represent individual cells; black lines show the mean, with light and dark grey areas marking 95% confidence intervals for the mean and standard deviation, respectively. Statistics: Z-test for two population proportions when comparing frequencies in (A), (B), (E); in univariate plot in (A) Kruskal-Wallis test with post hoc Dunn’s test; in (C) and (D, right): T-test for data that followed a normal distribution or Mann-Whitney U test otherwise; two-tailed t-test in (D, left). Symbols: **, P ≤ 0.01. Anaph., anaphase; org., organoids; m., middle; arb., arbitrary; cents, centrosomes; divis., division.

**Supplementary Figure 4.**
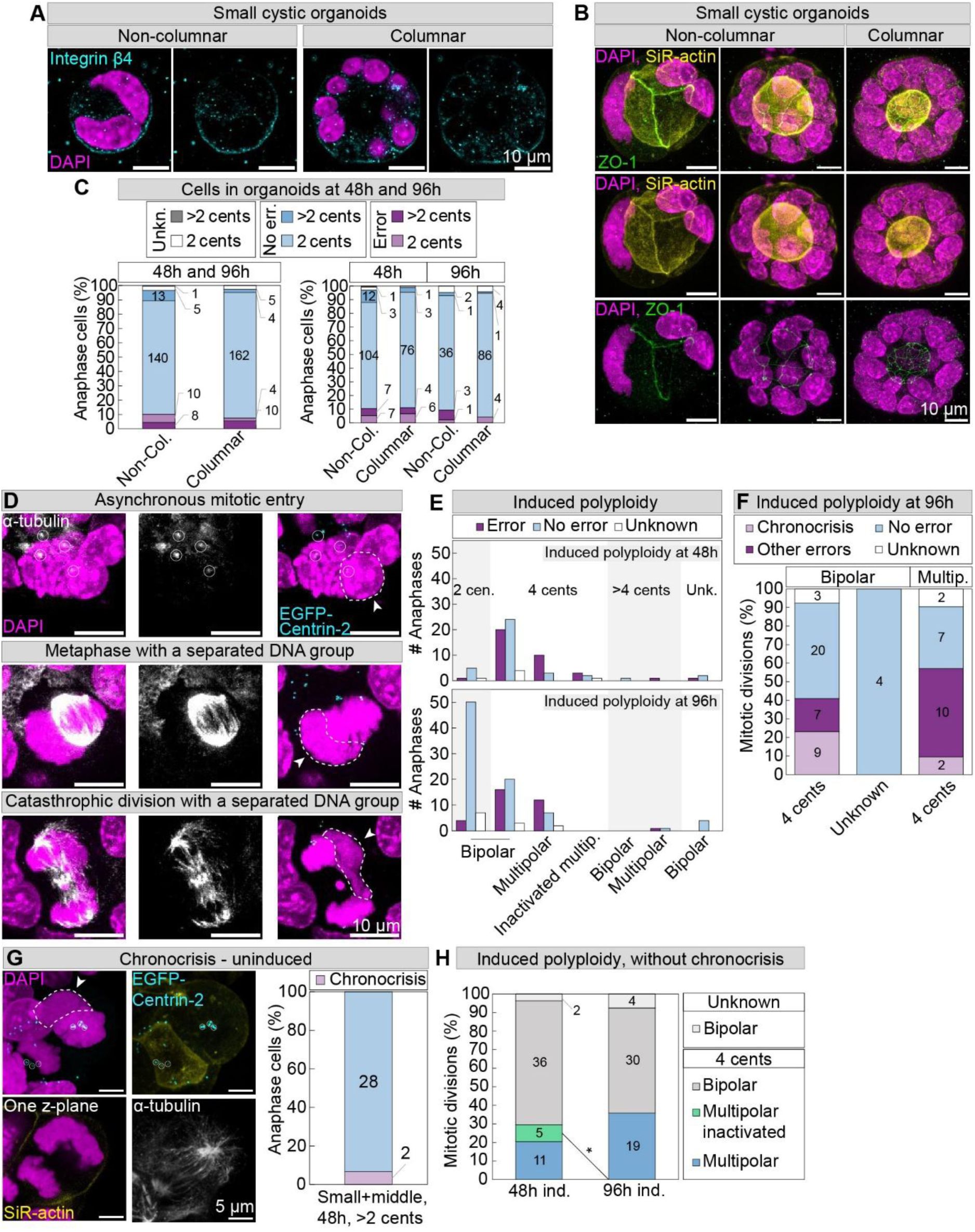
Early mSI organoids establish apicobasal polarity and often undergo error-prone chronocrisis after polyploidy induction. **(A)** Images show fixed mSI organoids immunostained for basal marker β4-integrin (cyan) and **(B)** apical marker Zonula occludens-1 (ZO-1, green) and stained with SiR-actin (yellow) and DAPI (magenta). Images that show β4-integrin staining are single z-planes. **(C)** The bar plot on the left shows the frequencies of chromosome segregation errors in mSI anaphase cells expressing EGFP-Centrin-2 (stained as in Figure 1A), depending on whether they come from columnar or non-columnar organoids, examples of which are shown in (A) and (B) under “Columnar” or “Non-columnar”. The bar plot on the right shows the same data, but with additional separation of cells by pseudotime of organoid growth, which is based on the time of fixation post seeding (48h and 96h) and the size of the organoid from which they were taken. Numbers of analysed cells are indicated on the data labels. Data from > 30 organoids and ≥ 3 experiments in each group. **(D)** Fixed images of mSI cells stably expressing EGFP-Centrin-2 (cyan), immunostained for α-tubulin (grey) and stained for F-actin with SiR-actin (not shown) and for DNA with DAPI (magenta), after polyploidy induction with cytochalasin D exhibiting: asynchronous mitotic entry of a binuclear cell in prophase (top row); separated chromatin groups of different condensation levels and lacking adjacent spindle poles in metaphase and anaphase, indicating chronocrisis (middle and bottom row). White arrows point to a nucleus of a binuclear that is late with mitotic entry, additionally encircled with a white dashed line (top row), or chromatin groups without adjacent poles, additionally encircled with white dashed lines (middle and bottom row). **(E)** The bar plots show the number of anaphase cells with denoted chromosome segregation outcomes after polyploidy induction at 48h (top) or 96h (bottom) after seeding dissociated organoids. Cells are separated based on their spindle polarity and centrosome numbers (color-coded shaded areas). Data from > 30 organoids and ≥ 3 experiments. **(F)** The bar plot shows the frequencies of chromosome segregation errors in anaphase cells after polyploidy induction at 96h post seeding dissociated organoids, separated by centrosome numbers and spindle polarity. Erroneous divisions coming from cells in chronocrisis are separated from other error types. Numbers of analysed cells are indicated on the data labels. Data from > 3 organoids and ≥ 3 experiments in each group. **(G)** Fixed mSI anaphase cell with a separated chromatin group without adjacent spindle poles, indicating chronocrisis. The bar plot on the right shows the frequency of anaphase cells in chronocrisis (purple) in mSI organoids 48h after seeding dissociated organoids. N = 30 from > 20 organoids and ≥ 3 experiments. **(H)** The bar plot shows frequencies of spindle polarity types in anaphase cells when chronocrisis events are excluded. Cells are separated based on polyploidy induction timepoint: at 48h and 96h after seeding dissociated organoids. Only cells with four centrosomes are shown, unless the number of centrosomes is unknown. N = (54, 53) cells from > 50 organoids and ≥ 3 experiments in each group. All images are maximum intensity projections, unless stated otherwise. “Uninduced” stands for native mSI organoids in which no polyploidy induction was performed, contrary to “induced polyploidy”. White circles mark centriole pairs for which centrosome identity was confirmed using α-tubulin staining. Statistics: Z-test for two population proportions in (A), (F), (H). Symbols: *, P < 0.05. Cents, centrosomes; unkn., unknown; no err., no error; non-col., non-columnar; col., columnar; multip., multipolar; ind., induced.

**Supplementary Figure 5.**
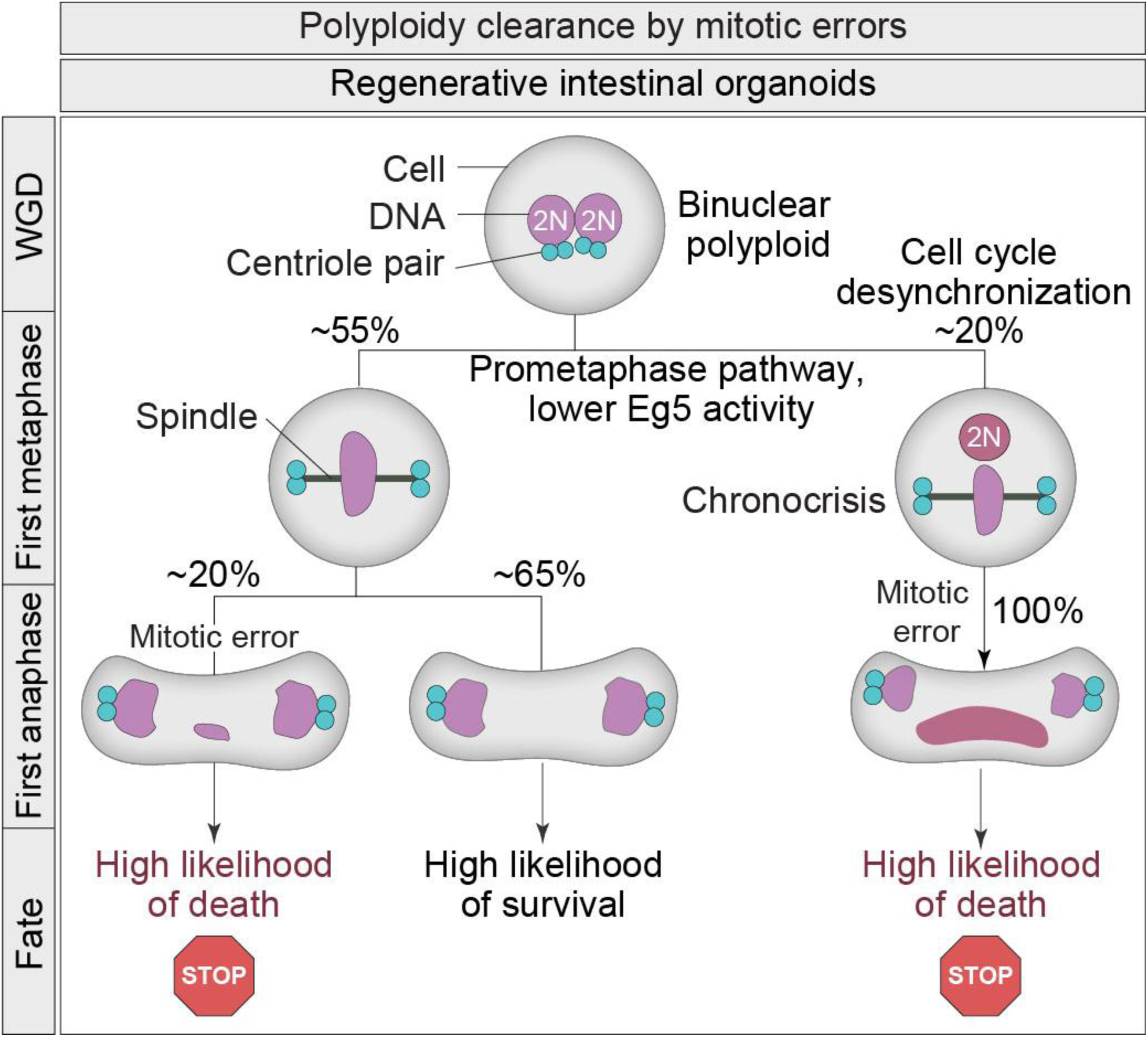
Transient global polyploidization impairs the formation of structured organoids. **(A)** The bar plot shows the frequencies of chromosome segregation errors in the first division of live-imaged mSI cells after polyploidy induction with cytochalasin D at 48h after seeding dissociated organoids. Cells are further separated based on spindle polarity. N = (33, 11) cells from > 3 organoids and ≥ 3 experiments. Representative live-imaged polyploid multipolar (top) and bipolar (bottom) erroneous anaphase of mSI cells stably expressing H2B-mNeon (color coded by depth and smoothed with 0.5 sigma Gaussian blur). **(B)** Large-field images of fixed mSI organoids stably expressing EGFP-Centrin-2 (not shown), stained for F-actin with SiR-actin (red) and for DNA with DAPI (blue). Organoids were either treated with DMSO for control (top) or cytochalasin D to induce polyploidy (bottom), 48h after seeding dissociated organoids. Time of fixation is 48h after DMSO or inhibitor wash-out. Images in corners are enlargements of representative organoid groups marked with white rectangles. All images are maximum intensity projections. Statistics: Z-test for two population proportions in (A). Symbols: **, P ≤ 0.01.

